# Structure-guided engineering of biased-agonism in the human niacin receptor via single amino acid substitution

**DOI:** 10.1101/2023.07.03.547505

**Authors:** Manish K. Yadav, Parishmita Sarma, Manisankar Ganguly, Sudha Mishra, Jagannath Maharana, Nashrah Zaidi, Annu Dalal, Vinay Singh, Sayantan Saha, Gargi Mahajan, Saloni Sharma, Mohamed Chami, Ramanuj Banerjee, Arun K. Shukla

**Author notes:** ^#^Joint 1^st^ author (Manish K. Yadav, Parishmita Sarma).

## Abstract

The Hydroxycarboxylic acid receptor 2 (HCA2), also known as the niacin receptor or GPR109A, is a prototypical G protein-coupled receptor that plays a central role in the inhibition of lipolytic and atherogenic activities in our body. Interestingly, GPR109A activation also results in vasodilation that is linked to the side-effect of flushing associated with dyslipidemia drugs such as niacin. This receptor continues to be a key target for developing novel pharmacophores and lead compounds as potential therapeutics in dyslipidemia with minimized flushing response, however, the lack of structural insights into agonist-binding and receptor activation has limited the efforts. Here, we present five different cryo-EM structures of the GPR109A-G-protein complexes with the receptor bound to dyslipidemia drugs, niacin or acipimox, non-flushing agonists, MK6892 or GSK256073, and recently approved psoriasis drug, monomethyl fumarate (MMF). These structures allow us to visualize the binding mechanism of agonists with a conserved molecular interaction network, and elucidate the previously lacking molecular basis of receptor activation and transducer-coupling. Importantly, cellular pharmacology experiments, guided by the structural framework determined here, elucidate pathway-selective biased signaling elicited by the non-flushing agonists. Finally, taking lead from the structural insights, we successfully engineered receptor mutants via single amino acid substitutions that either fail to elicit agonist-induced transducer-coupling or exhibits G-protein signaling bias. Taken together, our study provides previously lacking structural framework to understand the agonist-binding and activation of GPR109A, and opens up the possibilities of structure-guided novel drug discovery targeting this therapeutically important receptor.

## Introduction

The Hydroxycarboxylic acid receptor 2 (HCA2), also known as the niacin receptor or GPR109A, belongs to the superfamily of G protein-coupled receptors (GPCRs), and it is expressed primarily in the adipose tissues^1–3^, keratinocytes^4^, immune cells such as neutrophils and Langerhans cells^5^ in our body. Upon activation by agonists, GPR109A couples to Gαi sub-family of heterotrimeric G-proteins leading to lowering of cAMP response^6–9^. In addition, activated GPR109A also recruits β- arrestins^10^, which are multifunctional proteins involved in GPCR desensitization, trafficking and downstream signaling^11^. Interestingly, GPR109A was identified as the molecular target for the action of nicotinic acid (aka, niacin), an effective drug prescribed for lowering the triglycerides, almost two decades ago^12^. Moreover, GPR109A activation also mediates the lowering of LDL (aka, bad cholesterol), enhancing the levels of HDL (aka, good cholesterol)^13^. Furthermore, monomethyl fumarate (MMF), the active metabolite of a psoriasis drug, Fumaderm, and also a therapeutic agent for the treatment of relapsing forms of multiple sclerosis, has also been identified as an agonist of GPR109A^14–16^. However, activation of GPR109A is also responsible for driving the troublesome side effect of flushing response associated with niacin, acipimox and MMF^10,13,16–19^. This represents a potential limitation with their therapeutic usage, and therefore, additional small molecule agonists targeting GPR109A remains a key focus area^7,19–21^.

Several non-flushing agonists, such as MK6892 and GSK256073 with high affinity for GPR109A have been developed and characterized using *in-vitro* and animal studies although none of these compounds is yet approved for clinical usage^22,23^. In addition, a comprehensive study has also demonstrated that the side effect of niacin-induced flushing response in mouse is driven primarily by β-arrestin-mediated downstream signaling, and therefore, G-protein-biased agonists of GPR109A may represent improved therapeutics compared to niacin^10^ (**Figure 1A**). In the same study, a previously developed agonist MK0354 was reported to maintain the anti-lipolytic effect with significant reduction in flushing response, and it was further characterized as a G-protein-biased agonist^10^. Still however, direct structural visualization and molecular mechanism of agonist-binding and activation of GPR109A remain primarily elusive and represent an important knowledge gap to efficiently target this receptor for therapeutic benefits.

**Figure 1:**
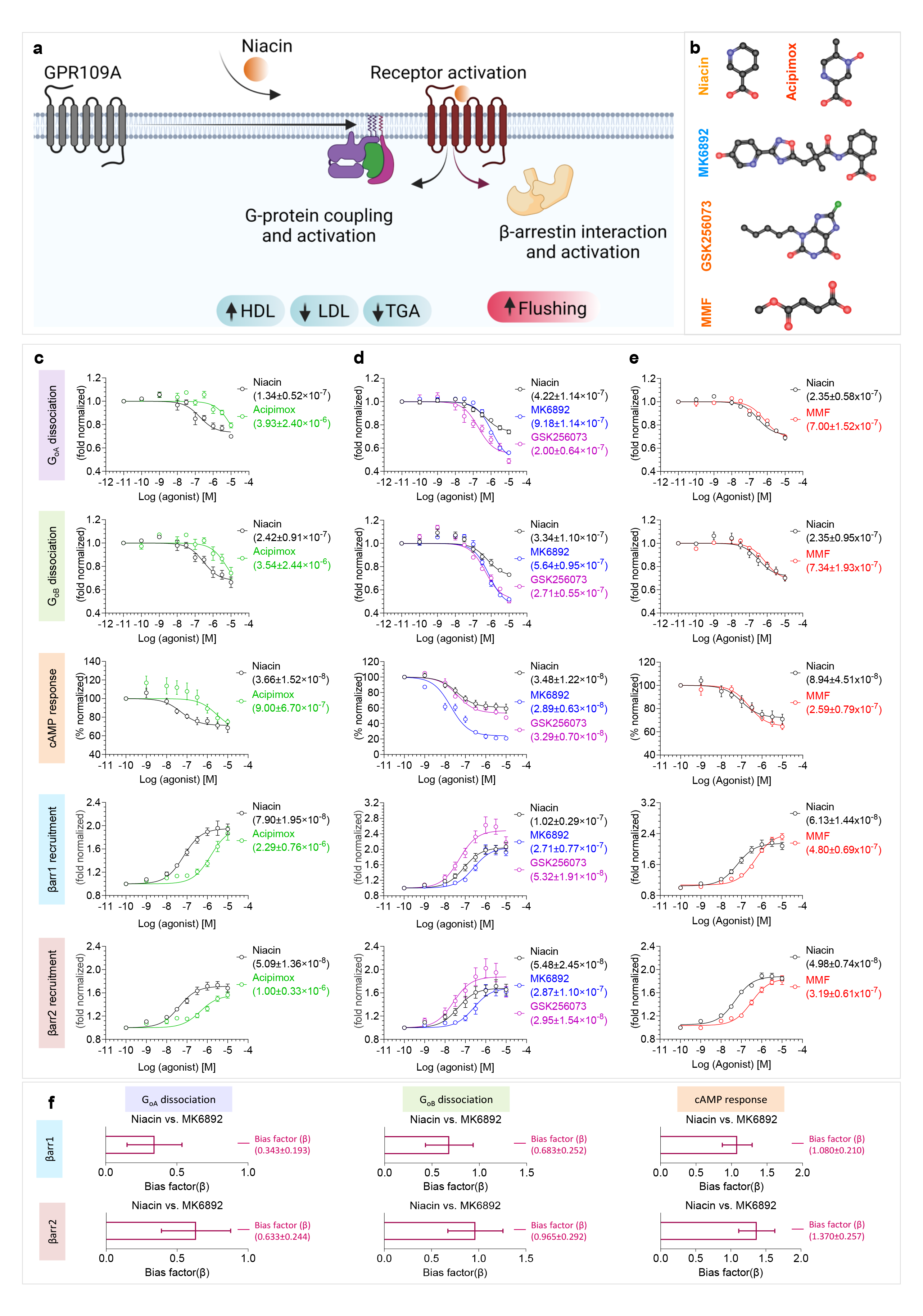
Pharmacological profiling of niacin, acipimox, MK6892, GSK256073 and MMF on GPR109A. **a,** Chemical structure of four GPR109A ligands used in the current study**. b,** Diagrammatic illustration of GPR109A activation and downstream signalling outcomes. **c,** G-protein activation and βarr recruitment downstream of GPR109A in response to acipimox with niacin as a reference ligand. First and second panel showing G_oA_ and G_oB_ dissociation studied by nanoBiT-based G-protein dissociation assay (Receptor+LgBiT-Gα_oA_/Gα_oB_+Gβ+SmBiT-Gγ) (mean±SEM; n=4; fold normalized with the minimum concentration for each ligand as 1). Forskolin induced cAMP decrease measured by GloSensor assay is shown in the third panel (mean±SEM; n=4; % normalized with the minimum concentration for each ligand as 100). βarr recruitment was studied by nanoBiT-based assay (Receptor-SmBiT+LgBiT-βarr) and is shown in fourth and fifth panel (mean±SEM; n=4 and n=5 for fourth and fifth panel respectively; fold normalized with the minimum concentration for each ligand as 1). **d,** G-protein activation and βarr recruitment downstream of GPR109A in response to MK6892 and GSK256073 with niacin as a reference ligand. First and second panel showing G_oA_ and G_oB_ dissociation (mean±SEM; n=4; fold normalized with the minimum concentration for each ligand as 1). Forskolin induced cAMP response is shown in the third panel (mean±SEM; n=4; % normalized with the minimum concentration for each ligand as 100). Fourth and fifth panel showing βarr1 and 2 recruitment respectively (mean±SEM; n=6; fold normalized with the minimum concentration for each ligand as 1). **e,** G-protein activation and βarr recruitment downstream of GPR109A in response to monomethyl fumarate (MMF) with niacin as a reference ligand. First and second panel showing G_oA_ and G_oB_ dissociation (mean±SEM; n=3; fold normalized with the minimum concentration for each ligand as 1). Forskolin induced cAMP response is shown in the third panel (mean±SEM; n=4; % normalized with the minimum concentration for each ligand as 100). Fourth and fifth panel showing βarr1 and 2 recruitment respectively (mean±SEM; n=4; fold normalized with the minimum concentration for each ligand as 1). **f,** Bias factor was calculated using the software https://biasedcalculator.shinyapps.io/calc/. During bias factor calculation Niacin stimulated response was considered as reference and observed G-protein biased with MK6892.

Here, we present five different cryo-EM structures of GPR109A-G-protein complexes where the receptor is activated either by niacin, acipimox, MK6892, GSK256073 and MMF. Comparison of these structural snapshots provides the molecular basis of ligand recognition, activation, and transducer-coupling by GPR109A. Importantly, the structural insights allow us to rationally design receptor mutants harbouring single amino acid substitution that either render the receptor completely inactive with respect to transducer-coupling, or, impart significant transducer-coupling bias. Our study not only illuminates the structural pharmacology of GPR109A ligands and paves the way for structure-guided discovery of novel therapeutics but also offers a framework to leverage the structural information to rationally encode signaling-bias in GPCRs.

## Results

### GPR109A agonists used for structural analysis

In order to visualize the molecular framework of ligand recognition and receptor activation, we selected five different ligands namely, niacin, acipimox, MK6892, GSK256073 and MMF (**Figure 1B**). Of these, niacin and acipimox are clinically prescribed drugs to treat dyslipidemia, while MK6892 and GSK256073 have been developed as non-flushing agonists of GPR109A. MK6892 is a biaryl cyclohexene carboxylic acid derivative that was reported to exhibit high affinity for GPR109A without significant off-target profile, and also displayed reduced vasodilation in animal studies while maintaining free fatty acid reduction similar to niacin^22^. GSK256073 was reported to display robust specificity for GPR109A over the other hydroxycarboxylic acid receptor subtypes, maintain the ability to lower the levels of non-esterified fatty acids in pre-clinical animal studies with reduced flushing response, and even promising outcomes in healthy male subjects^23^. MMF on the other hand, is the active metabolite of psoriasis drug, Fumaderm, and is also used as a therapeutic agent in multiple sclerosis^14,15,24^. Our selection of these ligands was based on their diverse chemical structures, therapeutic profile, and associated side effects with the goal to understand their interaction with GPR109A and potentially link the structural insights with their therapeutic profile.

We measured the pharmacological profile of acipimox, MK6892, GSK250673 and MMF with niacin as a reference agonist of GPR109A in G-protein response and β-arrestin recruitment assays (**Figure 1C-E**). We observed that acipimox behaved as a full agonist but with lower potency in G-protein dissociation, cAMP response, and β-arrestin recruitment assay (**Figure 1C**). Moreover, MK6892 exhibited higher efficacy in G-protein response and similar efficacy but lower potency in β- arrestin recruitment compared to niacin (**Figure 1D**). In the case of GSK256073, we observed a higher response in G-protein dissociation but similar efficacy in cAMP response, and it also displayed higher efficacy in β-arrestin recruitment as compared to niacin (**Figure 1D**). On the other hand, MMF behaved as a full agonist in both G-protein and β-arrestin recruitment assay with similar efficacy as niacin but slightly weaker potency in β-arrestin assays (**Figure 1E**). Analysis of these pharmacology data and calculation of the bias factor suggest that MK6892 acts as a G-protein biased agonist at GPR109A (**Figure 1F**). In these cellular assays, the surface expression of GPR109A was measured using a previously described whole cell-based ELISA method with mock-transfected cells as negative control (**Supplementary Figure 1).**

### Overall structure of agonist-GPR109A-G-protein complexes

We reconstituted the agonist-GPR109A-G-protein complexes using purified components following state-of-the-art methodology successfully applied to other GPCR-G-protein complexes (REF). We determined the cryo-EM structures of these complexes at estimated resolutions of 3.37 Å, 3.45 Å, 3.45 Å, 3.26 Å and 3.56 Å respectively, for the niacin, acipimox, MK6892, GSK256073 and MMF-bound receptor (**Figure 2A-E, Supplementary Figure 2-10**). The unambiguous densities of the coulombic maps enabled us to assign nearly the entire transmembrane domain of the receptor although the first seven residues at the N-terminus of the receptor and the last forty-five residues at the carboxyl-terminus were not resolved in the structures potentially due to their inherent flexibility (**Supplementary Figure 11**). Still however, in each of these complexes, the ligand densities were clearly discernible, allowing us to visualize ligand-receptor interactions, and the high map quality at the receptor-G-protein interface facilitated the identification of residue level interactions driving G-protein coupling to the receptor (**Supplementary Figure 12**). The precise sequence of the components resolved in these structures is listed in **Supplemental Figure 11**. The overall structures of GPR109A in all five complexes are highly similar with an RMSD of 0.6-1.0 Å^2^ along the Cα of the receptor interface (**Figure 3A**) and the key differences are observed in the ligand-receptor interaction as outlined in the sections below.

**Figure 2:**
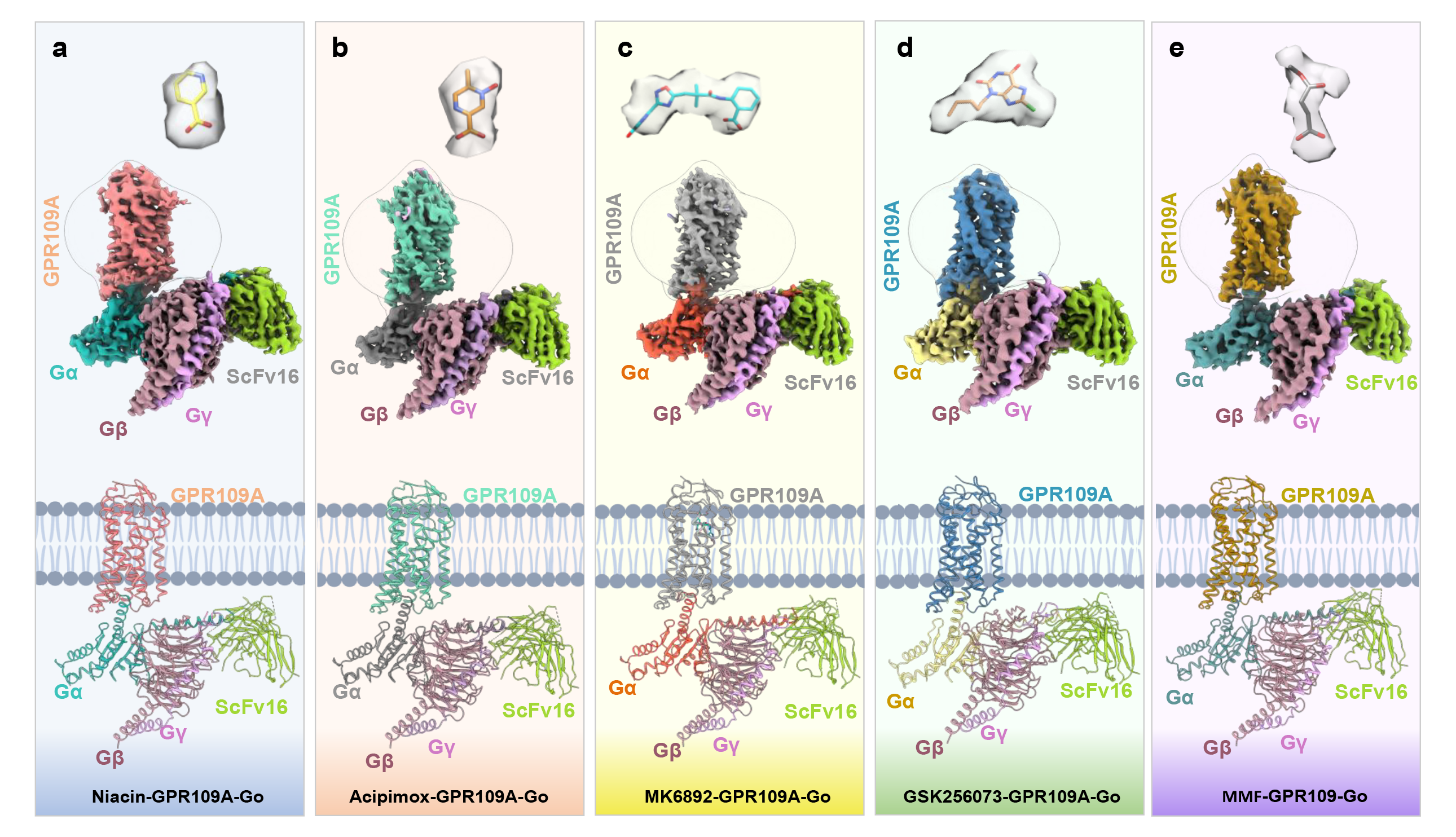
Overall architecture of Niacin, Acipimox, MK6892, GSK256073 and MMF bound GPR109A-G protein complexe. Map and ribbon diagram of the ligand-bound GPR109A-Go complexes (front view) and the cryo-EM densities of the ligands (sticks) are depicted as transparent surface representations. **a**, **niacin-GPR109A-Go**: Light coral: GPR109A, light sea green: miniGαo, rosy brown: Gβ1, orchid: Gɣ2, yellow green: ScFv16, **b**, **acipimox-GPR109A-Go**: medium aquamarine: GPR109A, gray: miniGαo, rosy brown: Gβ1, orchid: Gɣ2, yellow green: ScFv16, **c, MK6892-GPR109A-Go**: Dark gray: GPR109A, tomato: miniGαo, rosy brown: Gβ1, orchid: Gɣ2, yellow green: ScFv16, **d, GSK256073-GPR109A-Go**: Steel blue: GPR109A, khaki: miniGαo, rosy brown:Gβ1, orchid: Gɣ2, yellow green: ScFv16, **e**, **MMF-GPR109A-Go:** Dark golden rod: GPR109A, cadet blue: miniGαo, rosy brown:Gβ1, orchid: Gɣ2, yellow green: ScFv16.

**Figure 3:**
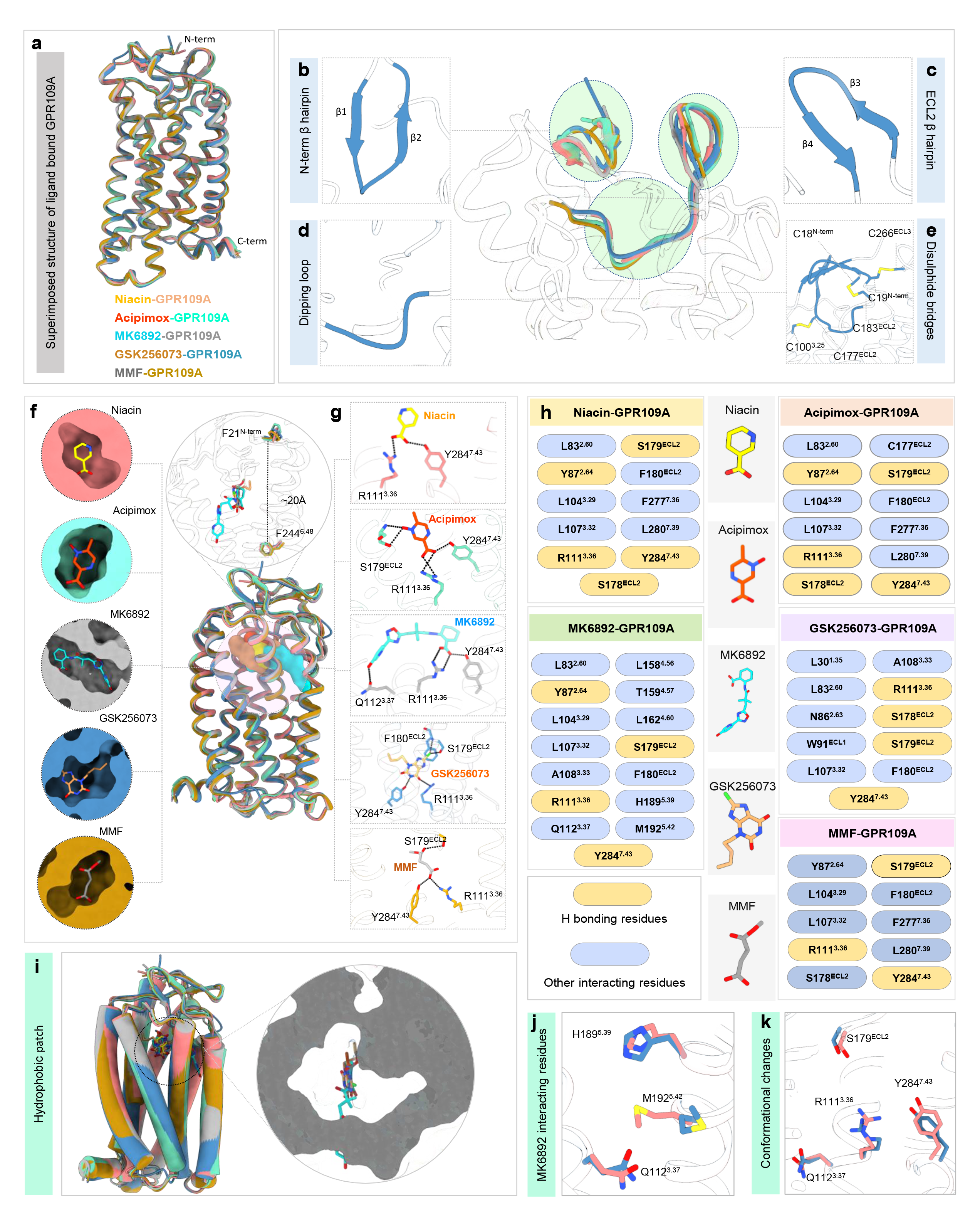
Ligand binding pocket of GPR109A. **a,** Structure superposition of niacin, acipimox, MK6892, GSK256073 and MMF bound GPR109A. **b,** Structural features of Ligand(GSK256073) bound-GPR109A, N terminal β-hairpin (left upper panel), **c,** Close-up view of ECL2 dipping into the orthosteric pocket (left lower panel), **d,** ECL2 β-hairpin (upper right panel) and **e,** Ribbon diagram of disulfide bridges. **f,** Superposed niacin, acipimox, MK6892, GSK256073 and MMF bound GPR109A structures highlighting the orthosteric binding pocket (Left panel, cross-sections of GPR109A bound to the individual ligand. **g,** GPR109A ligand binding pocket highlighting the major interactions of the individual ligand. **h,** List of GPR109A residues interaction with ligands. **i,** Cross-section of GPR109A orthosteric sites depicting the hydrophobic patch surrounding the individual ligand. **j,** Interacting residues of GPR109A with the extended part of MK6892. **k,** Conformational changes of GPR109A residues interacting with MK6892 with respect to niacin.

### An extracellular lid in GPR109A

Structural analysis of the receptor component in these structures uncovered several interesting features. For example, the N-terminus of the receptor in all five structures adopts a twisted β- hairpin structure that positions itself above the extracellular opening of the receptor **(Figure 3B)**. Moreover, the Ile169-Leu176 segment in the ECL2 adopts a twisted antiparallel β-hairpin confor-mation while Ser178^4.51^-Ser181^5.31^ dips down into the core of the TM bundle to form part of the or-thosteric binding pocket **(Figure 3C, D)**. Interestingly, the ECL2 hairpin interacts with the β-hairpin formed by the N-terminal residues Leu11-Cys18 to form a “lid-like” architecture that covers the ex-tracellular opening of the receptor **(Figure 3E)**. This observation can be attributed to the presence of three disulfide bridges (N-terminus Cys18 with Cys183 of ECL2, N-terminus Cys19 with Cys266 of TM7, and Cys177 of ECL2 with Cys100 of TM3) which helps to further stabilize the N-terminus-ECL2 lid **(Figure 3E)**. These disulfide bonds might impose additional constraints towards the flexi-bility of the lid, and facilitate docking of the ligand within the orthosteric pocket of the receptor. It is interesting to note that a similar “lid-like” conformation adopted by the N-terminus and ECL2 has been previously reported for several GPCRs such as b2AR and CXCR4, removal of the disulfide constraints resulting in either complete loss or decreased agonist affinity in the receptors (REF).

### Agonist-receptor interaction in the orthosteric binding pocket

The ligand binding site in GPR109A is positioned approximately 20 Å deep in the receptor core (measured from N-terminal Phe21 to Phe244^6.48^) (**Figure 3F**), and all five agonists share a common interaction interface, at least in part, on the receptor where chemically similar moieties of the ligands are positioned (**Figure 3F**). An array of aromatic residues namely, Phe276^7.35^, Phe277^7.36^, Trp91^ECL1^, Phe180^ECL2^ and Phe193^5.43^; and hydrophobic residues namely, Leu83^2.60^, Leu104^3.29^ and Leu107^3.32^ are found lining the orthosteric pocket of the receptor, and together, they form the microenvironment for the binding of the ligands. The interactions between niacin, acipimox, MK6892, GSK256073 and MMF with the receptor are mainly ionic, hydrophobic, and aromatic, including residues predominantly from TM2, TM3, TM7 and ECL2, and a complete list of interactions are listed in **Supplementary Figure 12**.

The comparison of the ligand binding pocket in all five structures reveals that Arg111^3.36^ forms the most important residue for binding to the negatively charged carboxyl group of ligands, niacin, acipimox, GSK256073, MK6892 and MMF through hydrogen bond (**Figure 3G**). Previous studies have suggested that this carboxyl group is critical for receptor activation, and substitution with an amide group abolishes GPR109A activity, and our structural snapshots provide a mechanistic basis for these functional observations (REF). Three more pairs of hydrogen bonds can be observed, one between the carboxyl moiety of niacin (or acipimox, GSK256073 and MMF) with the side chain Tyr284^7.43^, two between the chloride moiety of GSK256073 with the side chain Ser179^45.52^ and backbone N-atom of Phe180^ECL2^ (**Figure 3G**) and one between oxo-group at position 4 of MMF with Ser179^45.52^. Furthermore, activation of GPR109A appears to require a hydrophobic environment within the orthosteric pocket, and several hydrophobic contacts can be found to stabilize niacin and acipimox within the ligand binding pocket mediated by hydrophobic residues namely, Leu83^2.60^, Leu104^3.29^, Leu107^3.32^, Phe180^ECL2^, Phe277^3.36^, and Leu280^7.39^ (**Figure 3I and Supplementary Figure 12)**. Similarly, hydrophobic residues such as Leu30^1.35^, Trp91^ECL1^, Leu107^3.32^, and Phe180^ECL2^ forms extensive interactions with GSK256073 (**Figure 3I**). Although niacin, acipimox and GSK256073 exhibit hydrophilic, hydrophobic, and charged properties that largely match with those of the ligand binding pocket (**Supplementary Figure 12**), slight conformational variation can be observed within the binding pocket for the niacin or acipimox and GSK256073 complexes. These conformational shifts can be attributed to the presence of the extra Cl moiety in GSK256073.

Like niacin, acipimox and GSK256073, the carboxyl group of MK6892 makes similar contacts with the surrounding polar and hydrophobic residues within the orthosteric binding pocket (**Figure 3G, H**). MK6892 has a relatively extended chemical structure compared to the other three agonists and therefore, it engages several additional residues in the receptor. For example, the extended moieties in MK6892 i.e., dimethyl, oxadiazole, and pyridyl groups interact with Gln112^3.37^, His189^5.39^ and Met192^5.42^ in an extended binding pocket in the receptor (**Figure 3J**). Interestingly, several conformational rearrangements in the side-chains of Arg111^3.36^, Gln112^3.37^, Ser179^45.52^, and Tyr284^7.43^ are also observed compared to the other agonists in order to accommodate the bulky extended group of MK6892 (**Figure 3K**). Furthermore, an upward rotameric transition of His189^5.39^ and an outward shift of Met192^5.42^ is also observed within the extended binding pocket to prevent steric clashes with the extended chain of MK6892 (**Figure 3J)**. These additional interactions of MK6892 with the GPR109A are similar to those observed in a recent study^25,26^.

### Agonist-induced activation of GPR109A

When compared to the recently determined inactive state crystal structure of GPR109A, the niacin-activated GPR109A displayed the known conformational changes i.e., the cytoplasmic side of TM6 exhibits an outward movement of ∼4 Å (measured from the Cα of Lys227) and 5.5 Å inward movement of TM5 towards the extracellular side (measured from the Cα of His189^5.39^) and about 4.5 Å outward movement towards the cytoplasmic side (measured from the Cα of Arg218^ICL3^) (**Figure 4A-C**). The agonist-bound structures of GPR109A exhibit the typical hallmark movements of receptor activation as reflected by the conserved motifs and microswitches. For example, the “P-I-F motif” consisting of Pro200^5.50^, Ile115^3.40^ and Phe240^6.44^ forms an interface at the base of the ligand binding pocket, and it undergoes conformational rearrangements upon receptor activation. The rearrangements include: (i) rotameric shift of Pro200^5.50^, (ii) rotameric flip of Ile115^3.40^ and (iii) large transition of Phe240^6.44^, thus opening the cytoplasmic core of the receptor for the interaction with the α5 residues of Gαo (**Figure 4D**). Similar conformational changes can be observed with respect to the “D-R-Y” and “NPxxY” microswitches as well. Asp124^3.49^, Arg125^3.50^ and Tyr126^3.51^ in TM3 is a highly conserved motif where Asp124^3.49^ forms a salt bridge with Arg125^3.50^, thus locking the receptor in an inactive conformation. A rotameric shift of Arg125^3.50^ can be observed in the ligand-bound structures, facilitating the breaking of the salt-bridge/ionic lock and transition to its active conformation (**Figure 4D**). A variant of the “NPxxY” motif is present in GPR109A, where Asn290^7.49^ is substituted with Asp290^7.49^ in TM7. Upon activation, the lower portion of TM7 moves inwards towards the receptor core combined with a rotation of Tyr294^7.53^ along the helical axis (**Figure 4D)**.

**Figure 4.**
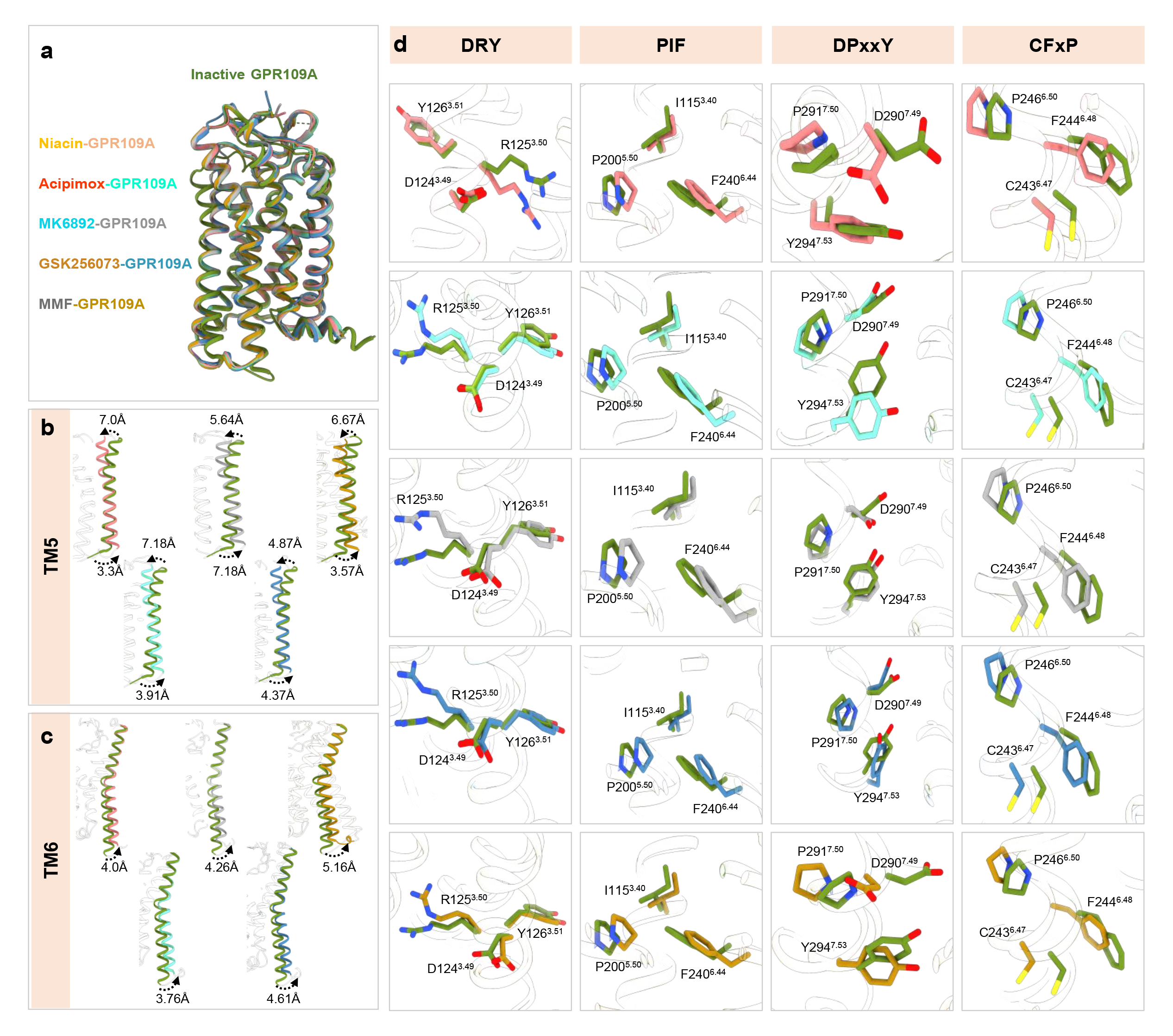
Major conformational changes on GPR109A activation. **a,** Superimposition of inactive GPR109A with receptor bound to niacin, acipimox, MK6892, GSK256073, and MMF. **b, c,** Displacements of TM5, TM6 upon GPR109A activation in the structures of niacin, acipimox, MK6892, GSK256073, and MMF bound GPR109A respectively. **d,** Conformational changes in the conserved microswitches (DRY, PIF, N/DPxxY, CW/FxP) in the active structure of GPR109A.

### The interface of GPR109A-G-protein interaction

As mentioned earlier, significant movements of TM5, TM6 and TM7 create an opening on the cytoplasmic surface of the receptor that allows the docking of the α5 helix of Gαo leading to coupling of G-proteins with the activated receptor (**Figure 5A, C, E, G, I**). Expectedly, we observe a large buried surface area at the interface of the receptor and G-protein nearing almost 2000 Å^2^ as typically observed in GPCR-G-protein complexes, and this is almost identical in all five structures of GPR109A reported here (**Figure 5A, C, E, G, I**). The GPR109A-G-protein interface is stabilized by extensive hydrophobic and polar interactions between the TM2, TM3, ICL2, ICL3, TM6, TM7 and H8 in the receptor and the α5 helix of Gαo (**Figure 5B, D, F, H, J**). Specifically, Tyr354 at the carboxyl-terminus of Gαo forms a key residue that is positioned in pocket on the cytoplasmic side of the receptor lined by Lys225^6.25^, Ile226^6.30^ and Pro299^8.48^ (**Figure 5B)**. In addition, several hydrogen bonds between Asp341, Asn347 and Gly352 of Gαo with Arg218^ICL3^, Arg128^3.53^ and Ser297^7.56^ of the GPR109A, respectively, further stabilize the interaction (**Figure 5B, D, F, H**). Finally, the stretch from Ala135 to Lys138 in the ICL2 of the receptor adopts a one-turn helix where His133 interacts with Thr340 and Ile342 which lie within a hydrophobic pocket formed by the residues from α5 helix, αN-β1 loop and β2-β3 loop of Gαo (**Figure 5F**). The receptor-G protein interface is further stabilized by residues of ICL3 with α5 C-terminal loop and α4-β6 loop of Gαo, viz. Arg218^ICL3^ forms extensive interactions with Thr340, Asp341 and Ile344 of Gαo (**Figure 5B, D, F, H**). A list of ligands bound-GPR109A residues interacting with Gαo is presented in **Figure 5K-L**, which underscores a largely conserved interface for G-protein interaction although some ligand-specific interactions are also observed. A comprehensive detail of the interactions between GPR019A and G-proteins are listed in **Supplementary Figure S13**.

**Figure 5:**
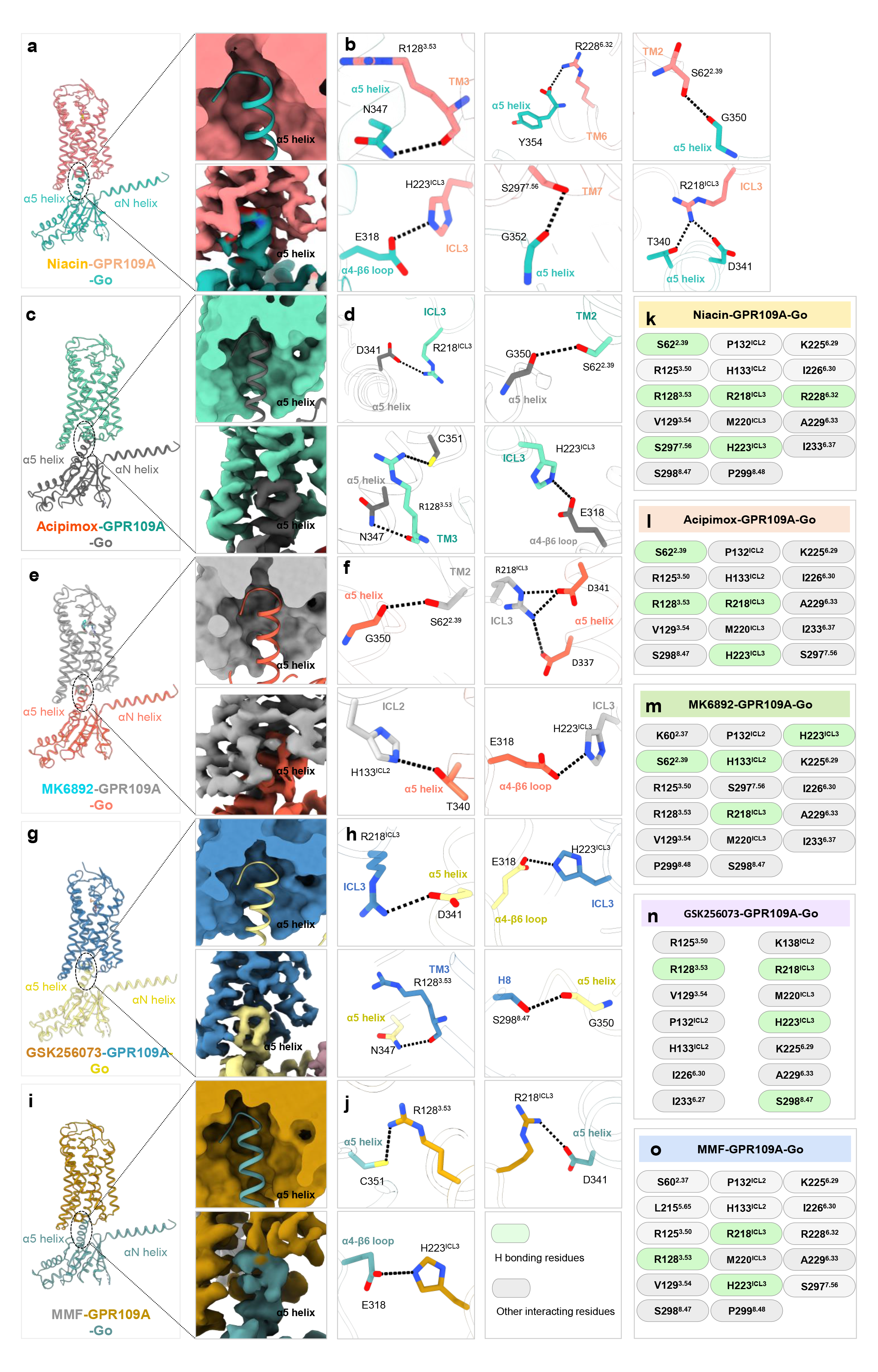
GPR109A-G-protein interacting interface. **a, c, e, g, i,** Representation of α5 helix of Gαo docking into the cytoplasmic core of GPR109A bound to niacin, acipimox, MK6892, GSK256073 and monomethyl fumarate (MMF) respectively. **b, d, f, h, j,** Key interactions between Gαo residues and residues of the cytoplasmic core of GPR109A. Black dotted line represents the H-bond. **k-o,** Illustration of residues contact between GPR109A and Gαo in niacin, acipimox, MK6892, GSK256073, and monomethyl fumarate (MMF) bound structures.

### Structure-guided design of receptor inactivation and biased-agonism

As mentioned above, there were two key interactions in the ligand binding pocket namely the Arg111 in TM3 and Ser179 in ECL2 that appeared to be conserved in all five structures involved in a direct hydrogen bonding with the ligands (**Figure 6A**). Therefore, we generated Arg111^3.36^Ala and Ser179^ECL2^Ala mutants of the receptor and measured niacin-induced G-protein and β-arrestin-coupling vis-à-vis the wild-type receptor. These mutants expressed at comparable levels as the wild-type receptor **(Supplementary Figure 14**). Interestingly, we observed that R111^3.36^A mutant exhibited complete loss of G-protein activation as measured using G-protein dissociation and cAMP assay, and agonist-induced β-arrestin recruitment (**Figure 6B-D**). This may reflect a near-complete loss of niacin binding to the receptor mutant as reported previously using a radioligand binding assay^2^. On the other hand, S179^ECL2^A mutation resulted in a significant reduction in β- arrestin recruitment in terms of Emax (1.61 fold vs. 1.47 fold for WT and S179^ECL2^A) and EC50 (32.0±4.90 nM vs. 3.05±65 nM for WT and S179^ECL2^A) (**Figure 6D**), while exhibiting slightly improved G-protein-coupling (Emax 38% vs. 53%, EC50 58.7±1.68 nM vs 9.90±1.42 nM for WT and S179^ECL2^A in Go dissociation assay, **Figure 6B**; Emax 25.78 vs. 64.95 in EC50 2.59±125 nM vs. 9.90±1.42 nM for WT and S179^ECL2^A in cAMP response assay, **Figure 6C**). Therefore, GPR109A^Ser179Ala^ mutant represents a G-protein-biased construct (**Figure 6E-F**), and it may be a useful tool to further probe ligand-bias at this receptor. Taken together, these data demonstrate the feasibility of structure-guided engineering of receptor inactivation and biased agonism at the transducer-coupling response, which may facilitate a framework to better understand the mechanistic aspects of biased agonism going forward.

**Figure 6.**
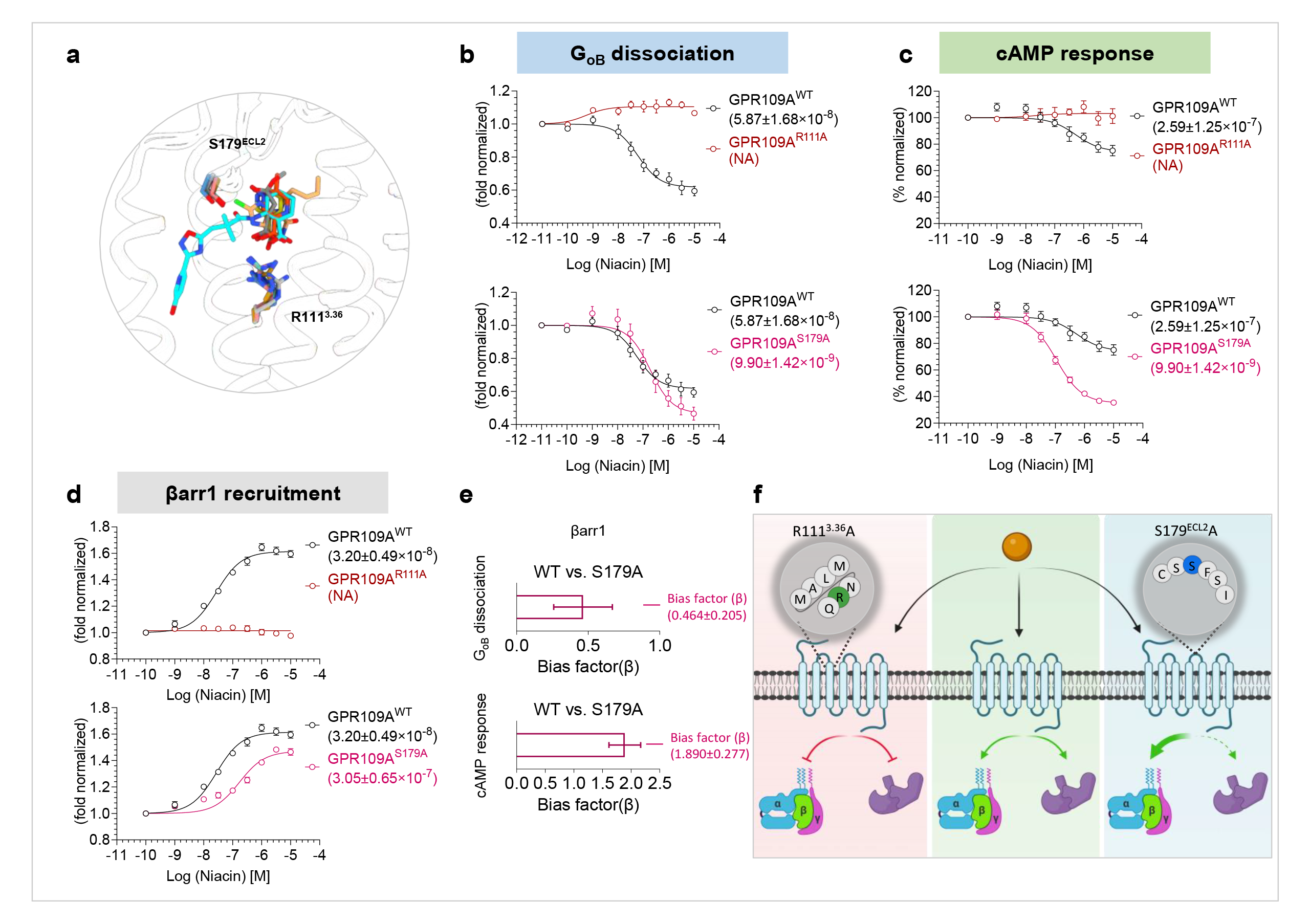
Structure guided bias signaling. **a,** Cartoon representation of residues interacting via H-bond with niacin (yellow), acipimox (orange red), MK6892 (cyan), GSK256073 (sandy brown), and monomethyl fumarate (MMF) (grey). **b,c,** G-protein activation downstream of GPR109A^WT^, GPR109A^R111A^, and GPR109A^S179A^ in response to niacin was studied by nanoBiT-based G-protein dissociation assay (Receptor+LgBiT-Gα_oB_+Gβ+SmBiT-Gγ) (panel b) (mean±SEM; n=3; fold normalized with the minimum concentration for each ligand as 1) and forskolin-induced cAMP level decay by GloSensor assay (panel c) (mean±SEM; n=3; % normalized with the minimum concentration for each ligand as 100). **d,** βarr recruitment downstream of GPR109A^WT^, GPR109A^R111A^, and GPR109A^S179A^ in response to niacin was studied by nanoBiT-based assay (Receptor-SmBiT+LgBiT-βarr) (mean±SEM; n=4; fold normalized with the minimum concentration for each ligand as 1). **e,** Bias factor for the mutant GPR109A^S179A^ was calculated using the software https://biasedcalculator.shinyapps.io/calc/. During bias factor calculation GPR109A^WT^ was considered as reference and observed G-protein biased with GPR109A^S179A^ upon stimulation with niacin. **f,** Schematic depicting the effect of the two mutants GPR109A^R111A^ and GPR109A^S179A^ on G-protein activation and βarr recruitment in response to niacin.

## Discussion

GPR109A continues to be an important drug target for developing therapeutic agents for dyslipidemia with properties superior to the commonly prescribed drug, niacin, especially in terms of reducing the side-effect of flushing response. While niacin, acipimox and MMF bind in a similar pose in the orthosteric binding pocket and make nearly-identical interactions, GSK256073 and MK6892 make additional contacts in the ligand binding pocket as expected based on their extended chemical structures. Interestingly, both GSK256073 and MK6892 appear to exhibit enhanced G-protein-coupling compared to niacin. However, GSK256073 is slightly more efficacious and potent in β-arrestin recruitment assay while MK6892 is slightly weaker than niacin in β-arrestin recruitment assay. Based on these data, it is tempting to speculate that additional contacts engaged by MK6892 make it a more potent and efficacious agonist of GPR109A compared to niacin, although follow-up experimentation would be required to test this hypothesis. As mentioned earlier, a previous study had reported the β-arrestin-mediated signaling to be the driver of flushing response for niacin while G-protein signaling is responsible for the lipid lowering effect. The G-protein-bias of MK6892 in β-arrestin recruitment assay therefore likely explains its non-flushing properties as reported earlier. However, the transducer-coupling profile of GSK256073 does not align with the same hypothesis, suggesting that the segregation of lipolysis vs. flushing response through GPR109A may involve additional fine-tuning that remains to be explored further.

Our mutagenesis studies guided by structural visualization of the key interactions between the agonists and the receptor yield a mutant that fails to elicit any transducer-coupling, and another mutant that maintains G-protein-coupling but loses β-arrestin binding. Although we have tested the effects of these mutations on only niacin-induced receptor activation and signaling, considering their conserved nature in terms of interaction with other agonists, it is likely that such mutants will exhibit a similar pattern for other agonists as well. It is intriguing that Ser179Ala mutation in ECL2 results in impaired β-arrestin recruitment as the site is closer to the orthosteric binding pocket in the receptor and significantly away from the interface of β-arrestin coupling to prototypical GPCRs. Therefore, it is tempting to speculate that the effect observed for Ser179^45.52^Ala mutation imparts a reduction in β-arrestin interaction through an allosteric mechanism as reported for other GPCRs previously^27,28^. In addition, the experimental framework established here should also facilitate the structure determination of other subtypes namely, HCA1 and HCA3, and a complete structural coverage of all three receptor subtypes should allow us to better understand the sub-type selectivity for niacin and other ligands.

In summary, the structural snapshots of GPR109A presented here elucidate the molecular details of the interaction of chemically-diverse agonists, and also uncover the mechanism of activation of this therapeutically important receptor. Our findings should pave the way for rational therapeutic design targeting GPR109A, and they also provide a framework to impart signaling bias in GPCRs guided by structural insights that may help deconvolute the mechanism of biased agonism going forward.

### Data availability statement

All the data are included in the manuscript and any additional information required to reanalyze the data reported in this paper is available from the corresponding author upon reasonable request.

## Acknowledgements

Research in A.K.S.’s laboratory is supported by the Senior Fellowship of the DBT Wellcome Trust India Alliance (IA/S/20/1/504916) awarded to A.K.S., Science and Engineering Research Board (IPA/2020/000405), Young Scientist Award from Lady Tata Memorial Trust, and IIT Kanpur. Cryo-EM was performed at BioEM lab of the Biozentrum at the University of Basel, and we thank Carola Alampi and David Kalbermatter for their excellent technical assistance.

## Authors’ contribution

MKY expressed and purified GPR109A, and reconstituted the receptor-G-protein complexes with help from VS and GM; PS carried out the pharmacological and cellular assays on GPR109A with help from SM, AD and NZ; SS purified mini-Gαo and ScFv16 with help from SaS, RB performed negative-staining EM, processed the cryo-EM data, prepared and deposited the coordinates with help from JM, and drafted the figures together with MG; MC screened the samples and collected cryo-EM data; AKS supervised and managed the overall project; all authors contributed to data analysis, interpretation and manuscript writing.

## Accession number

The cryo-EM maps and structures have been deposited in the EMDB and PDB with accession numbers 8IYP and EMD-35817 for niacin-GPR109A-Go, 8JER and EMD-36193 for acipimox-GPR109A-Go, 8IYW and EMD-35831 for GSK256073-GPR109A-Go, 8IYH and EMD-35822 for MK6892-GPR109A-Go and EMD-36280, PDB ID: 8JHN for MMF-GPR109A-Go complex.

## Materials and methods

### General reagents, plasmids, and cell culture

The majority of standard reagents were purchased from Sigma Aldrich unless mentioned. Dulbecco′s Modified Eagle′s Medium (DMEM), Phosphate Buffer Saline (PBS), Trypsin-EDTA, Fetal-Bovine Serum (FBS), Hank’s Balanced Salt Solution (HBSS), and Penicillin-Streptomycin solution were purchased from Thermo Fisher Scientific. HEK-293 cells were purchased from ATCC and maintained in 10% (v/v) FBS (Gibco, Cat. no. 10270-106) and 100U ml^-1^ penicillin and 100 µg ml^-1^ streptomycin (Gibco, Cat. no. 15140122) supplemented DMEM (Gibco, Cat. no. 12800-017) at 37 °C under 5% CO2. The cDNA coding region of GPR109A^WT^, GPR109A^Y87A^, GPR109A^R111A^, GPR109A^S179A^, and GPR109A^Y284A^ with a HA signal sequence, a FLAG tag followed by the N-terminal region of M4 receptor (2-23 residues) at the N-terminus was cloned into pcDNA3.1 vector. For GloSensor assay, luciferase-based 22F cAMP biosensor construct was purchased from Promega. For the constructs used in NanoBiT assay, SmBiT was fused at the C-terminus of the receptor, and the LgBiT-βarr1/2 construct was the same as previously described^29^. All DNA constructs were verified by sequencing from Macrogen. Niacin was purchased from Himedia (Cat. no. TC157), acipimox and MMF were purchased from Sigma Aldrich (Cat. no: 92571 and Cat. no: 651419), respectively. GSK256073 and MK9862 were purchased from MedChemExpress (Cat. no: HY10680 and HY119222, respectively).

### GPR109A purification

Codon-optimized human GPR109A was cloned in the pVL1393 vector with an N-terminal HA signal sequence followed by a FLAG tag and M4 receptor N-terminal sequence for increased expression. The receptor was expressed and purified from *Spodoptera frugiperda* (*Sf*9) cells using a baculovirus-mediated expression system. For receptor purification, insect cells were infected with recombinant baculovirus for 72 hrs at 27 °C and harvested by high-speed centrifugation. Post-harvest, insect cells were sequentially dounced in hypotonic buffer (20 mM HEPES, pH 7.4, 20 mM KCl, 10 mM MgCl2, 1 mM PMSF, and 2 mM Benzamidine), hypertonic buffer (20 mM HEPES, pH 7.4, 1 M NaCl, 20 mM KCl, 10 mM MgCl2, 1 mM PMSF, and 2 mM Benzamidine) and lysis buffer (20 mM HEPES, pH 7.4, 450 mM NaCl, 1 mM PMSF, and 2 mM Benzamidine). Lysed cells were solubilized by continuous stirring in 1% L-MNG (Anatrace, Cat. no. NG310) for two hours at 4 °C in the presence of 0.01% cholesteryl hemisuccinate (Sigma, Cat. no. C6512). To prevent receptor aggregation, 2 mM Iodoacetamide was added to the solution. Post-solubilization, salt concentration was lowered to 150 mM with dilution buffer (20 mM HEPES, pH 7.4, 2 mM CaCl2, 1 mM PMSF, and 2 mM Benzamidine), and cell debris was separated by high-speed centrifugation, and the receptor was enriched on M1-anti FLAG columns. Non-specific proteins were removed by three washes of low salt buffer (20 mM HEPES, pH 7.4, 150 mM NaCl, 2 mM EDTA, 0.01% MNG, 0.01% CHS, and 2 mM CaCl2) alternating with two washes of high salt buffer (20 mM HEPES, pH 7.4, 350 mM NaCl, 0.01% MNG, and 2 mM CaCl2). The bound receptor was eluted with FLAG peptide-containing buffer (20 mM HEPES, pH 7.4, 150 mM NaCl, 2 mM EDTA, and 250 µg ml^-1^ flag peptide). The purified receptor was treated with two rounds of 2 mM Iodoacetamide, followed by one round of 2 mM cysteine treatment to remove free Iodoacetamide. The purified receptor was concentrated using a 30 kDa MWCO concentrator (Cytiva, Cat no. 28932361) and stored at -80 °C with a 10% final glycerol concentration. 1 µM of niacin, acipimox, MK6892, GSK256073 or MMF was kept throughout the purification.

### Purification of Gβ1γ2 dimer

N terminal 6X His tagged Gβ1 and Gγ2 subunits were cloned in the Dual pVL1392 vector and expressed in the *Sf9* cell using a baculovirus-based expression system. For purification, insect cells were infected with the recombinant virus for 72 hrs and harvested with high-speed centrifugation. Cells were lysed by douncing in lysis buffer (50 mM Tris-Cl, pH 8.0, 150 mM NaCl, 1 mM PMSF, and 2 mM Benzamidine) and pelleted by centrifugation at 20,000 rpm for 20 min. The cell pellet was re-dissolved and dounced in solubilization buffer (50 mM Tris-Cl, pH 8.0, 150 mM NaCl, 5 mM β-mercaptoethanol, 1% DDM (Anatrace, Cat. no. D310), 1 mM PMSF, and 2 mM Benzamidine). Lysed cells were solubilized for two hrs at 4 °C with constant stirring. Cell debris was separated by high-speed centrifugation, and protein was passed through the Ni-NTA column using gravity flow. Non-specific proteins were removed by a one-column wash with buffer (50 mM Tris-Cl, pH 8.0, 150 mM NaCl, 0.01% MNG), and protein was eluted with 250 mM Imidazole (50 mM Tris-Cl, pH 8.0, 150 mM NaCl, 0.01% MNG, 250 mM Imidazole). Eluted protein was concentrated with a 10 kDa MWCO concentrator (Cytiva Cat. no. 28932360) and stored with 10% final glycerol concentration.

### Mini Gαo purification

The gene encoding miniG*α*o was designed as described previously^30,31^ and cloned into pET-15b (+) vector with 6X His tag at the N-terminal followed by TEV protease cleavage site. The recombinant construct was transformed into *E. coli* BL21(DE3) cells. A 5 ml starter culture, grown for 6-8h at 37 °C, was inoculated into a 50 ml primary culture media supplemented with 0.2% glucose and allowed to grow at 30 °C for 16-18 hrs. 1.5 litre of TB (Terrific Broth) media was inoculated with 15 ml of primary culture and grown at 30 °C. At O.D600 0.8, cells were induced with 50 μM IPTG (isopropylthio-β-galactoside) and allowed to grow for an additional 18–20 hrs. Cells were pelleted down and first lysed by lysozyme in lysis buffer (40 mM HEPES, pH 7.4, 100 mM NaCl, 10 mM Imidazole, 10% Glycerol, 5 mM MgCl2, 1 mM PMSF, 2 mM Benzamidine, 50 μM GDP, 100 μM DTT, and 1 mg ml^-1^ lysozyme), followed by disruption by ultrasonication. Cell debris was pelleted by high-speed centrifugation at 4 °C, and protein was enriched on the Ni-NTA column. Non-specifically bound proteins were removed by extensive washing with wash buffer (20 mM HEPES, pH 7.4, 500 mM NaCl, 40 mM Imidazole, 10% Glycerol, 50 μM GDP, and 1 mM MgCl2), and protein was eluted with 500 mM Imidazole (in 20 mM HEPES, pH 7.4, 100 mM NaCl, and 10% Glycerol). His tag was cleaved by overnight TEV treatment at room temperature (1:20, TEV: protein), and untagged protein was recovered by size exclusion chromatography on Hi-Load Superdex 200 PG 16/600 column (Cytiva, Cat. no. 17517501). Fractions corresponding to cleaved protein were pooled, analyzed on SDS-PAGE, and stored at -80 °C with 10% glycerol.

### ScFv16 purification

The gene encoding ScFv16 was cloned in pET-42a (+) vector downstream of 10X His tagged MBP gene with a TEV protease cleavage site between them and overexpressed in the *E. coli* Rosetta (DE3) strain^32^. A single colony from a freshly transformed plate was inoculated in 50 ml of 2XYT media and allowed to grow overnight at 37 °C. 1-litre 2XYT media supplemented with 0.5% glucose and 5 mM MgSO4 was inoculated with overnight primary culture and induced with 250 µM IPTG at O. D600 of 0.8–1.0. The culture was allowed to grow for 16–18 hrs at 18 °C. Post-harvest, cells were resuspended in 20 mM HEPES, pH 7.4, 200 mM NaCl, 30 mM Imidazole, 1 mM PMSF, and 2 mM Benzamidine buffer and incubated at 4 °C for 40 min with constant stirring. Cells were lysed by sonication, and cell debris was removed with high-speed centrifugation at 4 °C. Protein was captured on the Ni-NTA column using gravity flow, and non-specific proteins were removed by extensive washing with 20 mM HEPES, pH 7.4, 200 mM NaCl, and 50 mM Imidazole. Bound ScFv16 was eluted with 300 mM imidazole-containing buffer (20 mM HEPES, pH 7.4, 200 mM NaCl) and was re-passed through amylose resins, and after one column wash with 20 mM HEPES, pH 7.4, 200 mM NaCl buffer, bound protein was eluted with 10 mM maltose (prepared in 20 mM HEPES, pH 7.4, 200 mM NaCl). To obtain tag-free ScFv16, the eluted protein was overnight digested with TEV, and His-MBP was removed by passing the digested protein through the Ni-NTA column. Eluted protein was further cleaned by size exclusion chromatography on Hi-Load Superdex 200 PG 16/600 column. Eluted protein was analyzed on SDS-PAGE and stored at - 80 °C with 10% glycerol.

### Reconstitution of GPR109A-G protein-ScFv16 complexes

Purified GPR109A was mixed with a 1.2 molar excess of miniGo, Gβy, and ScFv16 in the presence of 25 mU ml^-1^ apyrase (NEB, Cat. no. M0398S) and 1 µM of individual ligand, and complexing was allowed for two hours at room temperature. The receptor complex was concentrated with a 100 kDa MWCO (Cytiva, Cat. no. GE28-9323-19) concentrator and separated from the unbound component by size exclusion chromatography on Superose 6 increase 10/300 GL column (Cytiva, Cat. no. 29091596). The SEC eluate was analyzed on 12% SDS-PAGE, and complex fractions were concentrated to ∼10 mg ml^-1^ and stored at -80 °C.

### Negative stain electron microscopy

Homogeneity of the purified protein complexes was determined through negative staining with uranyl formate prior to data collection under cryogenic conditions following the protocols described previously^33,34^. 3.5 µl of the purified complexes were dispensed onto fresh glow discharged carbon/formvar coated 300 mesh Cu grids (PELCO, *Ted Pella*) at a concentration of 0.02 mg ml^-1^ and incubated for 1 min at room temperature. This was followed by blotting of the excess samples from the grids using filter paper. The grid containing the adhered sample was touched onto a first drop of freshly prepared 0.75% uranyl formate stain and immediately blotted off by touching the edge of the grid onto a filter paper. The grid was then touched and incubated on a second drop of uranyl formate for 30s and left on the bench in a Petri plate for air drying. Data collection was performed on a FEI Tecnai G2 12 Twin TEM (LaB6) operating at 120 kV and equipped with a Gatan CCD camera (4k x 4k) at 30,000x magnification. Data processing of the collected micrographs was performed with Relion^35^ 3.1.2 version. More than 10,000 particles were autopicked with the gaussian blob picker, extracted and subjected to reference-free 2D classification.

### Cryo-EM sample preparation and data collection

3 µl of the individual complexes were applied onto glow-discharged Quantifoil holey carbon grids (Cu R2/1 or R2/2) and vitrified in liquid ethane (−181 °C) using a Leica GP plunger (Leica Microsystems, Austria) maintained at 90% humidity and 10 °C. CryoEM movies were acquired on a TFS Glacios microscope operating at 200 kV and equipped with Gatan K3 direct electron detector (Gatan Inc.). Images were collected automatically with SerialEM software in counting mode at a nominal magnification of 46,000x and pixel size of 0.878 Å over a defocus range of 0.5-2.5 µm. An accumulated dose of 55 e^-^/A^2^ was fractionated into a movie stack consisting of 40 frames.

### Cryo-EM data processing

All data processing steps were performed with cryoSPARC^36^ v4.0 unless otherwise stated. Dose fractionated movie stacks were subjected to beam-induced motion correction using Patch motion correction (multi) followed by estimation of contrast transfer function parameters with Patch CTF estimation (multi).

For the Niacin-GPR109A-Go dataset, 11,070 dose weighted, motion-corrected micrographs were selected for downstream processing. Auto-picking yielded 7,027,107 particles which were subjected to several rounds of reference-free 2D classification to eliminate particles with poor features. 1,737,725 particle projections corresponding to the 2D averages with clear secondary features were selected and subjected to Ab-initio reconstruction with 3 classes. Subsequent heterogeneous refinement yielded a model with features of a typical GPCR-G protein complex containing 1,011,301 particle projections which accounted for 75% of the particles used for heterogeneous refinement. This particle stack was subjected to non-uniform refinement, followed by local refinement with mask excluding the noise outside the molecule, yielding a coulombic map with an indicated global resolution of 3.37 Å at 0.143 FSC cut-off.

For the MK6892-GPR109A-Go dataset, 90,683,101 particles were autopicked from 10,753 motion corrected micrographs which were extracted with a box size of 360 px (fourier cropped to 64 px) and subjected to multiple rounds of reference-free 2D classification. 2D class averages consisting of 1,829,840 particles with clear secondary features and resembling conformation of protein complexes were re-extracted with a box size of 360 px and fourier cropped to 288 px. These particle stacks from the extraction job were subsequently subjected to Ab-initio reconstruction, followed by heterogeneous refinement yielding 4 models. 1,200,513 particle projections (accounting for 66% of the total particles) from the best 3D class were selected and subjected to non-uniform refinement, followed by local refinement, which yielded a map with an overall resolution of 3.26 Å using the 0.143 FSC criterion.

For the GSK256073-GPR109A-Go dataset, 5,761,414 particles were automatically picked from 10,574 motion-corrected micrographs. These particle projections were extracted with a box size of 360 px (fourier cropped to 64 px) and subjected to iterative rounds of reference-free 2D classification to discard noisy particles. 1,601,694 particles corresponding to the 2D classes with evident features of protein complexes were selected and re-extracted with a box size of 360 px and fourier cropped to 288 px. These selected particle projections were used to generate 3 maps for heterogeneous refinement. One of the 3D classes with 523,816 particles showing all the features of a GPCR-G protein complex was subjected to 3D non-uniform refinement, reaching a nominal resolution of 3.45 Å.

For the Acipimox-GPR109A-Go dataset, 9,115,816 particles were autopicked from 11,263 micrographs, extracted with a box size of 360 px (fourier cropped to 64 px) followed by 2D classification to obtain classes with clear secondary features. 166,548 particles corresponding to the clean classes were re-extracted with a box size of 360 px (fourier cropped to 288) and subjected to ab-initio reconstruction and heterogenous refinement to generate 4 classes. 1,059,994 particles from the best 3D class were selected and subjected to non-uniform refinement followed by local refinement with mask to yield a final reconstruction at a resolution of 3.45 Å.

For the MMF-GPR109A-Go dataset, autopicking was performed with the blob-picker subprogram which yielded 9,029,435 particles. Particles were extracted with a box size of 360 px (fourier cropped to 64 px) and pared down to 183,241 particles after reference-free 2D classification. The clean particle stack was then re-extracted with a box size of 360 px (fourier cropped to 288 px). Two rounds of ab initio and subsequent hetero-refinement (using four models) were then performed to further refine the particle stack to 678,286 particles. Non-uniform refinement and successive local refinement resulted in a map with an estimated global resolution of 3.56 Å.

Local resolution estimation of all maps was determined using the Blocres subprogram within cryoSPARC with the corresponding half maps. Sharpening of all maps was performed with “Autosharpen maps” within the Phenix suite^37,38^ to enhance features for model building.

### Model building and refinement

Coordinates from an AlphaFold model of GPR109A (AF-Q8TDS4-F1) was used to dock into the EM density map of niacin-GPR109A-Go using Chimera^39,40^. Similarly, coordinates of Gαo, Gβγ and ScFv16 were obtained from a previously solved structure of C5aR1 in complex with Gαo (PDB: 8HPT). The combined model so obtained containing all the components was subjected to “all atom refine” sub-module within COOT^41^, followed by manual rebuilding of the residues and the ligand. The rebuilt model was subjected to real space refinement in Phenix^37,38^ to obtain a model with 97.13% of the residues in the most favoured region and 2.77% in the allowed region of the Ramachandran plot. Validation of all the models was performed with Molprobity^42^ within Phenix.

The ligand free model of niacin-GPR109A-Go complex (PDB: 8IY9) was fitted into the density maps of acipimox-GPR109A-Go and MMF-GPR109A-Go in Chimera followed by flexible fitting of the coordinates with the “all atom refine” module in COOT. After several rounds of manual adjustments, the generated model was automatically refined with Phenix_refine. The final refined models of acipimox-GPR109A-Go and MMF-GPR109A-Go showed good Ramachandran statistics with 96.86% and 97.31% in the most favored regions of the Ramachandran plot, respectively.

The ligand-free model of niacin-GPR109A-Go complex (PDB: 8IY9) was fitted into the density maps of acipimox-GPR109A-Go and MMF-GPR109A-Go in Chimera followed by flexible fitting of the coordinates with the “all atom refine” module in COOT. After several rounds of manual adjustments, the generated model was automatically refined with Phenix_refine. The final refined models of acipimox-GPR109A-Go and MMF-GPR109A-Go showed good Ramachandran statistics with 96.86% and 97.31% in the most favored regions of the Ramachandran plot, respectively.

Likewise, the ligand-free model of niacin-GPR109A-Go complex (PDB: 8IY9) was used to dock into the coulombic maps of MK6892-GPR109A-Go and GSK256073-GPR109A-Go using Chimera. The docked model and the corresponding maps were then imported into COOT and fitted into the respective maps with the “all atom refine” module. The poorly fitted regions were manually adjusted in COOT followed by iterative refinement of the coordinates against the maps using Phenix_refine. The final refined models of MK6892-GPR109A-Go and GSK256073-GPR109A-Go contained residues in 97.67% and 97.14% of the most favored regions of the Ramachandran plot with no outliers. Data collection, processing and model refinement statistics are provided in **Supplementary Figure 8**. All figures included in the manuscript have been prepared with Chimera and ChimeraX software.

### GloSensor-based cAMP assay

cAMP response upon ligand stimulation was measured by GloSensor assay^43^. Briefly, HEK-293 cells were transiently transfected with 2 μg of GPR109A construct together with 5 μg 22F cAMP plasmid using the transfection reagent polyethyleneimine (PEI) linear (Polysciences, Cat. no. 23966) at DNA: PEI ratio of 1:3. After 16-18hrs of transfection, cells were harvested followed by resuspension of the cell pellet in assay buffer composed of 1X HBSS, 20 mM of 4-(2- hydroxyethyl)-1-piperazineethanesulfonic acid (HEPES) pH 7.4 and D-luciferin (0.5 mg ml^-1^) (GoldBio, Cat. no.: LUCNA-1G). Harvested cells were then seeded in an opaque flat bottom white 96 well cell culture plate (SPL life sciences, Cat. no. 30196) at a density of 2×10^5^ cells well^-1^. After seeding, cells were incubated at 37 °C for 90 min and 30 min at room temperature. After 120 min of incubation basal level luminescence was recorded using a multi-mode plate reader (Lumistar/Fluostar microplate reader, BMG Labtech). In order to record ligand-induced cAMP decrease as a readout of Gi activation, cellular cAMP level was increased by adding 5 μM forskolin, and 7-8 cycles luminescence was recorded until the signal got saturated. Once the luminescence signal got stabilized cells were stimulated with corresponding ligands, and luminescence values were recorded for 20 cycles. For stimulation, ligand concentrations ranging from 100 pM to 10 μM were prepared by serial dilution in the buffer constituted of 1X HBSS, 20 mM HEPES pH 7.4. Nicotinic acid, acipimox, GSK256073, MK6892 and MMF of different concentrations were added to the corresponding wells. Baseline corrected data were normalized with respect to the luminescence signal of minimal concentration of each ligand as 100% and plotted using nonlinear regression analysis in GraphPad Prism v 9.5.0 software.

### Surface expression assay

Plasma membrane expression of receptors in respective assays was measured by whole cell-based surface ELISA as previously discussed^44^. Briefly, transfected cells were seeded at a density of 2×10^5^ cells well^-1^ in 0.01% poly-D-Lysine pre-treated 24-well plate and incubated for 24 h at 37 °C. Post incubation, growth media was aspirated, and cells were washed with ice-cold 1X TBS for once, followed by fixation with 4% PFA (w/v in 1X TBS) on ice for 20 min. Post fixation, cells were washed three times with 1X TBS (400 μl in each wash) followed by blocking with 1% BSA (w/v in 1X TBS) at room temperature for 90 min. After blocking with 1% BSA, 200 μl anti-FLAG M2-HRP was added and incubated for 90 min (prepared in 1% BSA, 1:10,000) (Sigma, Cat. no. A8592). Post antibody incubation, to remove unbound antibodies, cells were washed with 1% BSA (prepared in 1X TBS) three times, followed by the development of signal by treating cells with 200 μl TMB-ELISA (Thermo Scientific, Cat no. 34028) until the light blue colour appeared. Signal was quenched by transferring the light blue-coloured solution to a 96-well plate containing 100 μl 1M H2SO4. The absorbance of the signal was measured at 450 nm using a multi-mode plate reader. Next, cells were incubated with 0.2% Janus Green (Sigma; Cat. no. 201677) w/v for 15 min after removal of TMB-ELISA by washing once with 1X TBS. Afterwards, Janus Green was aspirated followed by washing with distilled water to remove the excess stain. After washing, 800 μl of 0.5 N HCl was added to elute the stain. 200 μl of the eluate was transferred to a 96-well plate, and at 595 nm absorbance was recorded. For analysis, data were analyzed by calculating the ratio of absorbance at 450/595 followed by normalizing the value of pcDNA transfected cells reading as 1. Normalized values were plotted using GraphPad Prism v 9.5.0 software.

### NanoBiT-based βarr recruitment assay

Plasma membrane localization of βarr upon stimulation of GPR109A with respective ligands was measured by luminescence-based enzyme-linked complement assay (NanoBiT-based assay) following the protocol described earlier^29,33,45^. Briefly, a receptor harbouring SmBiT at the carboxy-terminus (3.5 µg) and βarr1/2 constructs (3.5 µg) with N-terminally fused LgBiT were co-transfected in HEK-293 cells using the transfection reagent polyethyleneimine (PEI) linear at DNA: PEI ratio of 1:3. Post 16-18 hr of transfection, cells were trypsinized, and resuspended in the NanoBiT assay buffer containing 1X HBSS, 0.01% BSA, 5 mM HEPES pH 7.4, and 10 μM coelenterazine (GoldBio, Cat. no. CZ05). Cells were then seeded in opaque flat bottom white 96 well plate at a density of 1×10^5^ cells well^-1^ and incubated for 120 min (90 min at 37 °C, followed by 30 min at room temperature). Post incubation, basal level luminescence readings were taken, followed by ligand addition. A series of ligand concentrations, spanning from 10 pM to 10 μM, were prepared using a buffer solution composed of 1X HBSS and 5 mM HEPES at pH 7.4. Subsequently, cells were stimulated with different doses of the specified ligands. Luminescence upon stimulation was recorded up to 20 cycles by a multi-mode plate reader. For analysis, stimulated readings were normalized with respect to the signal of minimal ligand concentration as 1 and plotted using nonlinear regression analysis in GraphPad Prism v 9.5.0 software.

### NanoBiT-based G-protein dissociation assay

Agonist-induced G-protein activation was measured by a nanoBiT-based G-protein dissociation assay described previously^33^. Briefly, HEK-293 cells were transfected with 1 μg of LgBiT-tagged Gα subunit, 4 μg of SmBiT-tagged Gγ2 subunit, 4 μg of untagged Gβ1 subunit along with 1 μg of untagged receptor construct using transfection reagent PEI at DNA: PEI ratio of 1:3. Post transfection, cells were harvested and seeded in a 96 well plate at a density of 1×10^5^ cells well-1. Cells were seeded in buffer containing 1X HBSS, 0.01% BSA, 5 mM HEPES pH 7.4, and 10 μM coelenterazine and incubated for 120 min (90 min at 37 °C and 30 min at room temperature). Post incubation, 3 cycles of basal level luminescence readings were recorded using a multi-mode plate reader. After that, cells were stimulated with varying ligand concentrations ranging from 10 pM to 10 μM. After stimulation, 20 cycles of luminescence were recorded. For data analysis, values after 15 min of stimulation were used and normalized with respect to the signal at the minimal ligand concentration of 100%. Normalized values were plotted using nonlinear regression analysis in GraphPad Prism v 9.5.0 software.

## References

1. Soga, T. et al. Molecular identification of nicotinic acid receptor. Biochem Biophys Res Commun 303, 364–9 (2003).

2. Tunaru, S., Lattig, J., Kero, J., Krause, G. & Offermanns, S. Characterization of determinants of ligand binding to the nicotinic acid receptor GPR109A (HM74A/PUMA-G). Mol Pharmacol 68, 1271–80 (2005).

3. Wise, A. et al. Molecular identification of high and low affinity receptors for nicotinic acid. J Biol Chem 278, 9869–74 (2003).

4. Hanson, J. et al. Nicotinic acid-and monomethyl fumarate-induced flushing involves GPR109A expressed by keratinocytes and COX-2-dependent prostanoid formation in mice. J Clin Invest 120, 2910–9 (2010).

5. Maciejewski-Lenoir, D. et al. Langerhans cells release prostaglandin D-2 in response to nicotinic acid. Journal of Investigative Dermatology 126, 2637–2646 (2006).

6. Offermanns, S. et al. International Union of Basic and Clinical Pharmacology. LXXXII: Nomenclature and Classification of Hydroxy-carboxylic Acid Receptors (GPR81, GPR109A, and GPR109B). Pharmacol Rev 63, 269–90 (2011).

7. Blad, C.C., Ahmed, K., AP, I.J. & Offermanns, S. Biological and pharmacological roles of HCA receptors. Adv Pharmacol 62, 219–50 (2011).

8. Richman, J.G. et al. Nicotinic acid receptor agonists differentially activate downstream effectors. Journal of Biological Chemistry 282, 18028–18036 (2007).

9. Tang, Y.T. et al. Enhancement of arachidonic acid signaling pathway by nicotinic acid receptor HM74A. Biochemical and Biophysical Research Communications 345, 29–37 (2006).

10. Walters, R.W. et al. beta-Arrestin1 mediates nicotinic acid-induced flushing, but not its antilipolytic effect, in mice. J Clin Invest 119, 1312–21 (2009).

11. Reiter, E. & Lefkowitz, R.J. GRKs and beta-arrestins: roles in receptor silencing, trafficking and signaling. Trends Endocrinol Metab 17, 159–65 (2006).

12. Tunaru, S. et al. PUMA-G and HM74 are receptors for nicotinic acid and mediate its anti-lipolytic effect. Nat Med 9, 352–5 (2003).

13. Pike, N.B. Flushing out the role of GPR109A (HM74A) in the clinical efficacy of nicotinic acid. Journal of Clinical Investigation 115, 3400–3403 (2005).

14. Tang, H., Lu, J.Y.L., Zheng, X.M., Yang, Y.H. & Reagan, J.D. The psoriasis drug monomethylfumarate is a potent nicotinic acid receptor agonist. Biochemical and Biophysical Research Communications 375, 562–565 (2008).

15. Berger, A.A., et al. Monomethyl Fumarate (MMF, Bafiertam) for the Treatment of Relapsing Forms of Multiple Sclerosis (MS). Neurol Int 13, 207–223 (2021).

16. Hanson, J., Gille, A. & Offermanns, S. Role of HCA(2) (GPR109A) in nicotinic acid and fumaric acid ester-induced effects on the skin. Pharmacol Ther 136, 1–7 (2012).

17. Benyo, Z. et al. GPR109A (PUMA-G/HM74A) mediates nicotinic acid-induced flushing. J Clin Invest 115, 3634–40 (2005).

18. Benyo, Z., Gille, A., Bennett, C.L., Clausen, B.E. & Offermanns, S. Nicotinic acid-induced flushing is mediated by activation of epidermal langerhans cells. Mol Pharmacol 70, 1844–9 (2006).

19. Offermanns, S. Heating up the cutaneous flushing response. Arterioscler Thromb Vasc Biol 34, 1122–3 (2014).

20. Dubrall, D. et al. Do dimethyl fumarate and nicotinic acid elicit common, potentially HCA(2) -mediated adverse reactions? A combined epidemiological-experimental approach. Br J Clin Pharmacol 87, 3813–3824 (2021).

21. Bodor, E.T. & Offermanns, S. Nicotinic acid: an old drug with a promising future. Br J Pharmacol 153 **Suppl 1**, S68–75 (2008).

22. Shen, H.C. et al. Discovery of a Biaryl Cyclohexene Carboxylic Acid (MK-6892): A Potent and Selective High Affinity Niacin Receptor Full Agonist with Reduced Flushing Profiles in Animals as a Preclinical Candidate. Journal of Medicinal Chemistry 53, 2666–2670 (2010).

23. Sprecher, D. et al. Discovery and characterization of GSK256073, a non-flushing hydroxy-carboxylic acid receptor 2 (HCA2) agonist. European Journal of Pharmacology 756, 1–7 (2015).

24. van Veldhoven, J.P.D. et al. Structure-activity relationships of trans-substituted-propenoic acid derivatives on the nicotinic acid receptor HCA2 (GPR109A). Bioorganic & Medicinal Chemistry Letters 21, 2736–2739 (2011).

25. Yang, Y. et al. Structural insights into the human niacin receptor HCA2-G(i) signalling complex. Nat Commun 14, 1692 (2023).

26. Xin Pan, F.Y., Peiruo Ning, Zhiyi Zhang, Binghao Zhang, Geng Chen, Wei Gao, Chen Qiu, Zhangsong Wu, Kaizheng Gong, Jiancheng Li, Jiang Xia, Yang Du. Structural insights into ligand recognition and selectivity of the human hydroxycarboxylic acid receptor HCAR2. bioRxiv 2023.03.28.534513 (2023).

27. Ma, N., Nivedha, A.K. & Vaidehi, N. Allosteric communication regulates ligand-specific GPCR activity. FEBS J 288, 2502–2512 (2021).

28. Nivedha, A.K. et al. Identifying Functional Hotspot Residues for Biased Ligand Design in G-Protein-Coupled Receptors. Mol Pharmacol 93, 288–296 (2018).

29. Kawakami, K. et al. Heterotrimeric Gq proteins act as a switch for GRK5/6 selectivity underlying beta-arrestin transducer bias. Nat Commun 13, 487 (2022).

30. Carpenter, B. & Tate, C.G. Expression, Purification and Crystallisation of the Adenosine A(2A) Receptor Bound to an Engineered Mini G Protein. Bio Protoc 7(2017).

31. Nehme, R. et al. Mini-G proteins: Novel tools for studying GPCRs in their active conformation. PLoS One 12, e0175642 (2017).

32. Hong, C. et al. Structures of active-state orexin receptor 2 rationalize peptide and small-molecule agonist recognition and receptor activation. Nat Commun 12, 815 (2021).

33. Pandey, S. et al. Intrinsic bias at non-canonical, beta-arrestin-coupled seven transmembrane receptors. Mol Cell 81, 4605–4621 e11 (2021).

34. Ghosh, E. et al. Conformational Sensors and Domain Swapping Reveal Structural and Functional Differences between beta-Arrestin Isoforms. Cell Rep 28, 3287–3299 e6 (2019).

35. Zivanov, J. et al. New tools for automated high-resolution cryo-EM structure determination in RELION-3. Elife 7(2018).

36. Punjani, A., Rubinstein, J.L., Fleet, D.J. & Brubaker, M.A. cryoSPARC: algorithms for rapid unsupervised cryo-EM structure determination. Nat Methods 14, 290–296 (2017).

37. Liebschner, D. et al. Macromolecular structure determination using X-rays, neutrons and electrons: recent developments in Phenix. Acta Crystallogr D Struct Biol 75, 861–877 (2019).

38. Adams, P.D. et al. PHENIX: a comprehensive Python-based system for macromolecular structure solution. Acta Crystallogr D Biol Crystallogr 66, 213–21 (2010).

39. Pettersen, E.F. et al. UCSF ChimeraX: Structure visualization for researchers, educators, and developers. Protein Sci 30, 70–82 (2021).

40. Pettersen, E.F. et al. UCSF Chimera--a visualization system for exploratory research and analysis. J Comput Chem 25, 1605–12 (2004).

41. Emsley, P. & Cowtan, K. Coot: model-building tools for molecular graphics. Acta Crystallogr D Biol Crystallogr 60, 2126–32 (2004).

42. Chen, V.B. et al. MolProbity: all-atom structure validation for macromolecular crystallography. Acta Crystallogr D Biol Crystallogr 66, 12–21 (2010).

43. Baidya, M. et al. Allosteric modulation of GPCR-induced beta-arrestin trafficking and signaling by a synthetic intrabody. Nat Commun 13, 4634 (2022).

44. Pandey, S., Roy, D. & Shukla, A.K. Measuring surface expression and endocytosis of GPCRs using whole-cell ELISA. Methods Cell Biol 149, 131–140 (2019).

45. Maharana, J. et al. Structural snapshots uncover a key phosphorylation motif in GPCRs driving beta-arrestin activation. Mol Cell (2023).

