## Supplemental Material for "Structure-guided engineering of biased-agonism in the human niacin receptor via single amino acid substitution"

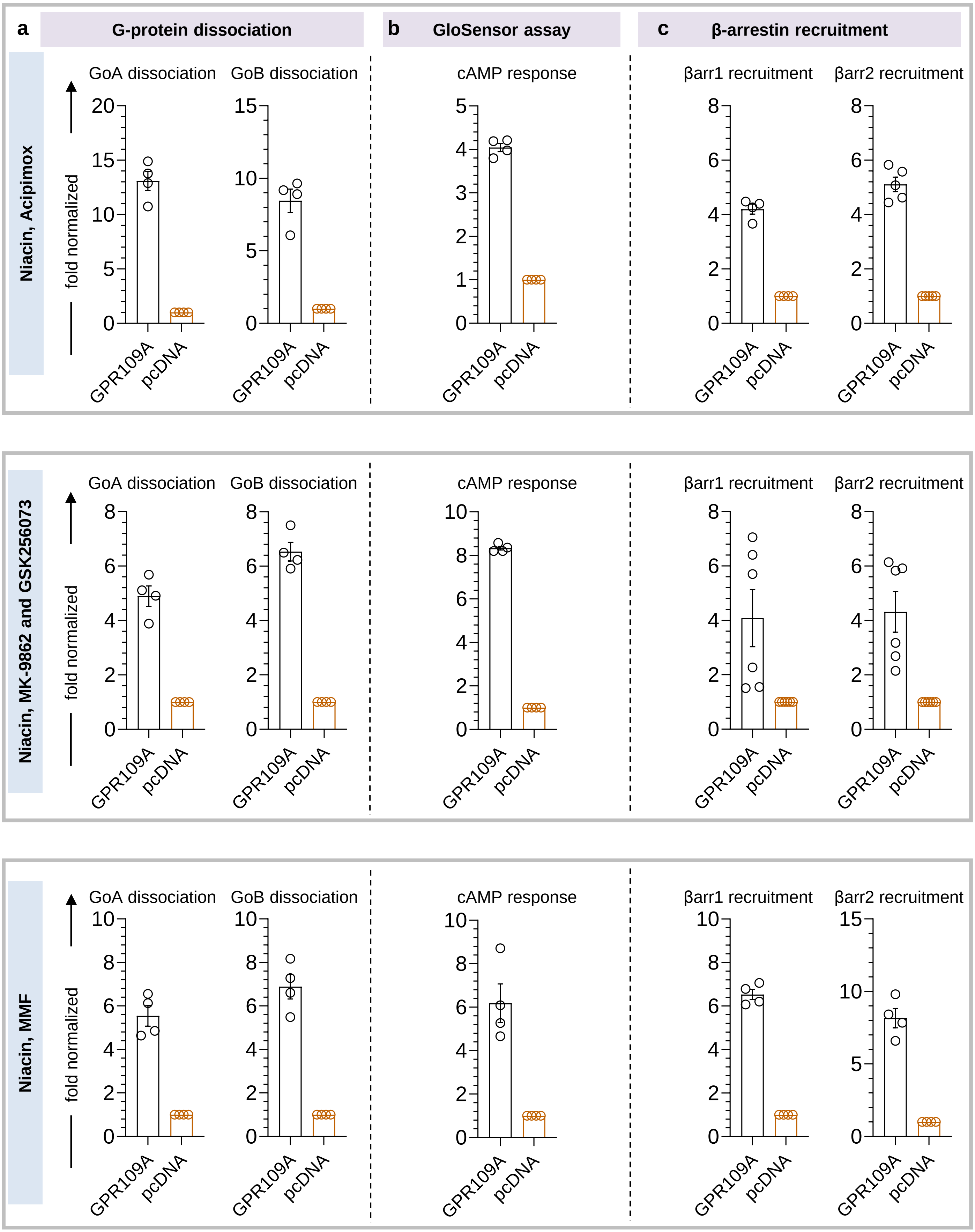


**Supplementary Figure 1: Surface expression of GPR109A in various assays.**

**a,** Surface expression of GPR109A in Go dissociation assay was measured using whole cell based surface ELISA (mean±SEM; n=4; normalized as fold over pcDNA) **b,** GPR109A surface expression in GloSensor assay (mean±SEM; n=4; normalized as fold over pcDNA) **c,** Surface expression of GPR109A in β-arrestin recruitment assay (mean±SEM; n=4 and n=5 for βarr1 and βarr2 recruitment respectively, upper panel, and n=6 for βarr1 and βarr2 recruitment; normalized as fold over pcDNA)

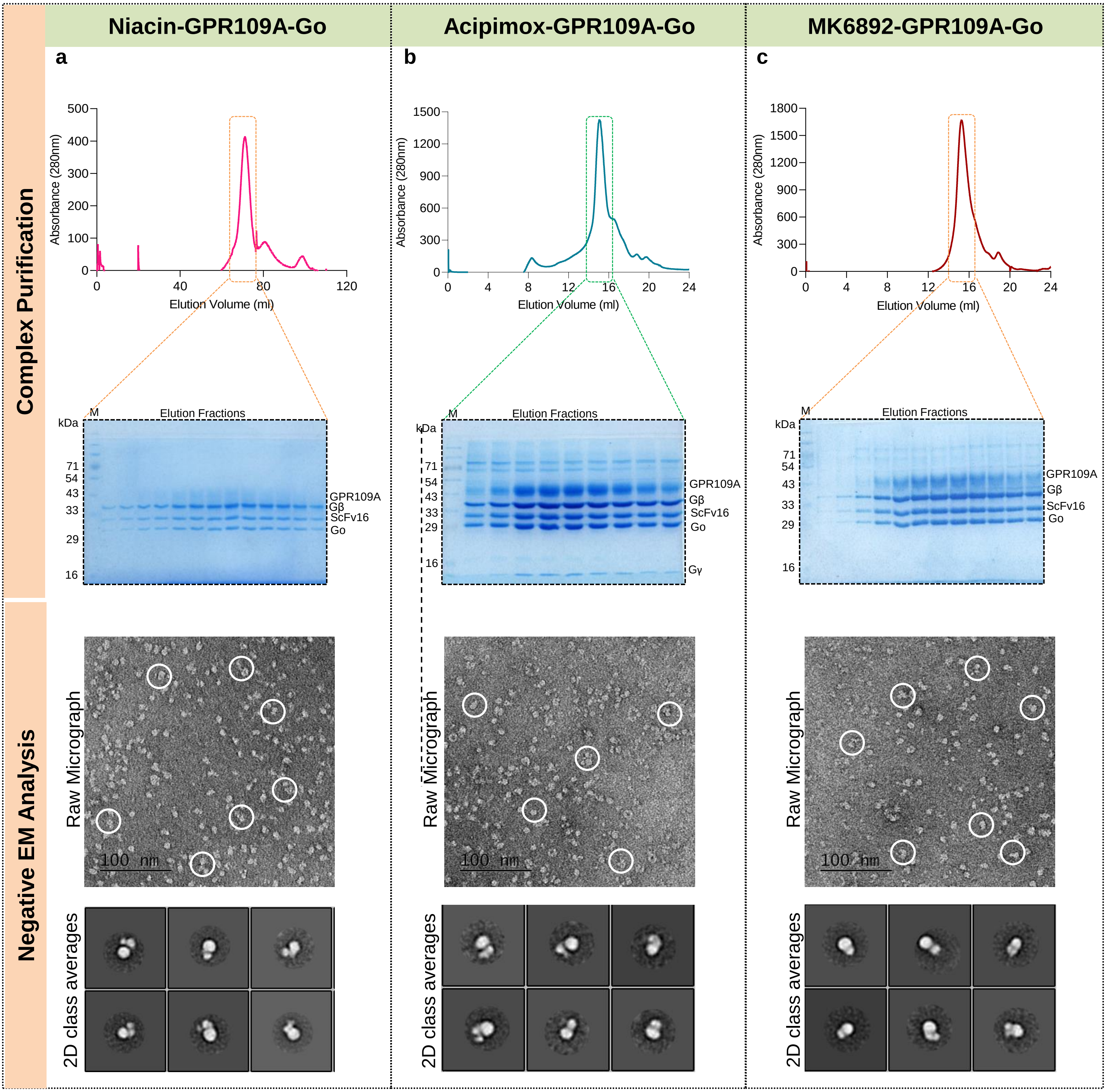


**Supplementary Figure 2. GPR109A-Go Complex Reconstitution and Visualization by Negative Staining EM. a, b, c.** Size exclusion chromatogram, SDS-PAGE analysis and negative stain EM analysis of niacin-GRP109A-Go, acipimox-GPR109A-Go, and MK6892-GPR109A-Go complexes, respectively.

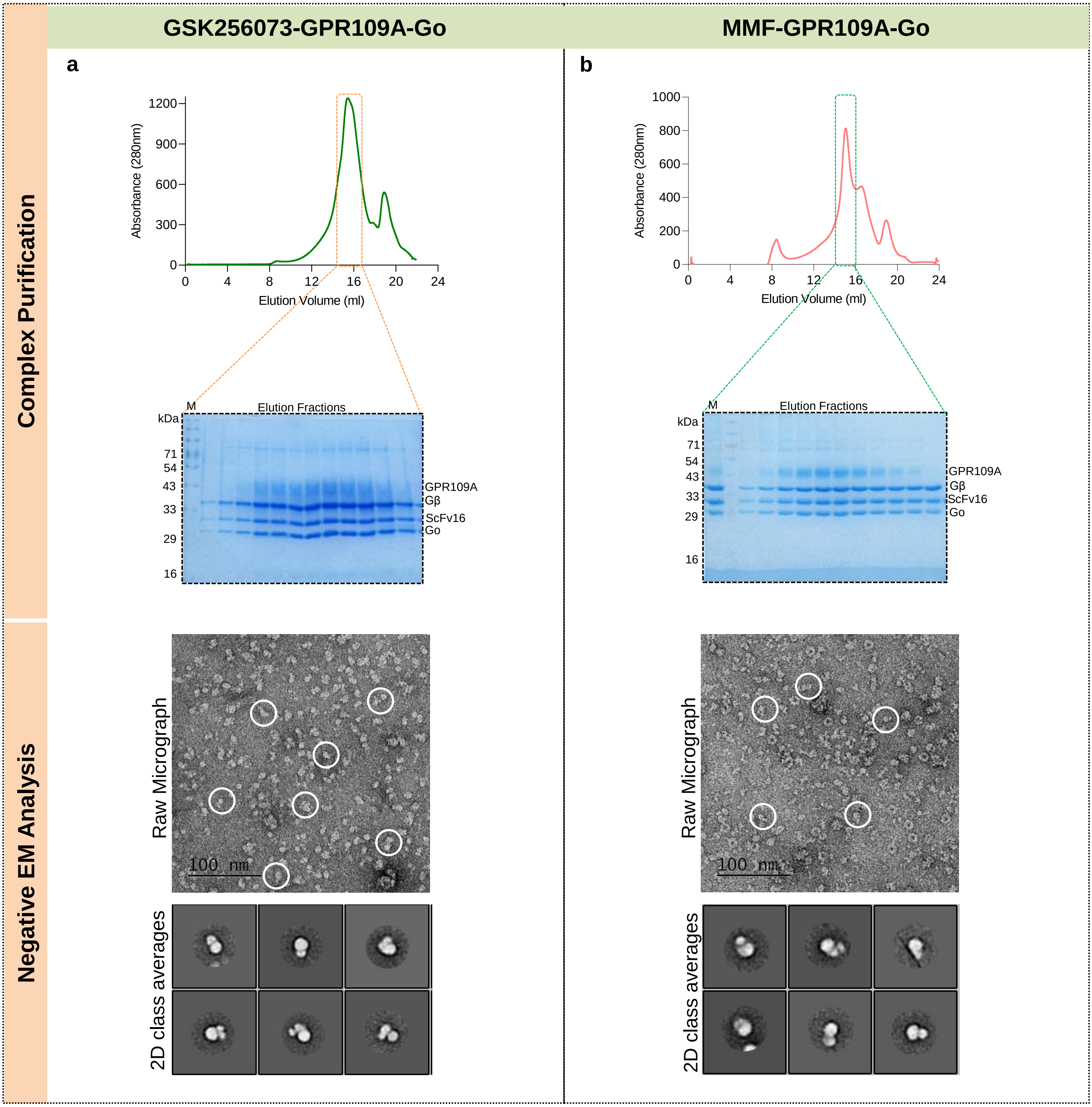


**Supplementary Figure 3. GPR109A-Go Complex Reconstitution and Visualization by Negative Staining EM. a, b**. Size exclusion chromatogram, SDS-PAGE analysis and negative stain EM analysis of GSK256073-GRP109A-Go and MMF-GPR109A-Go complex, respectively.

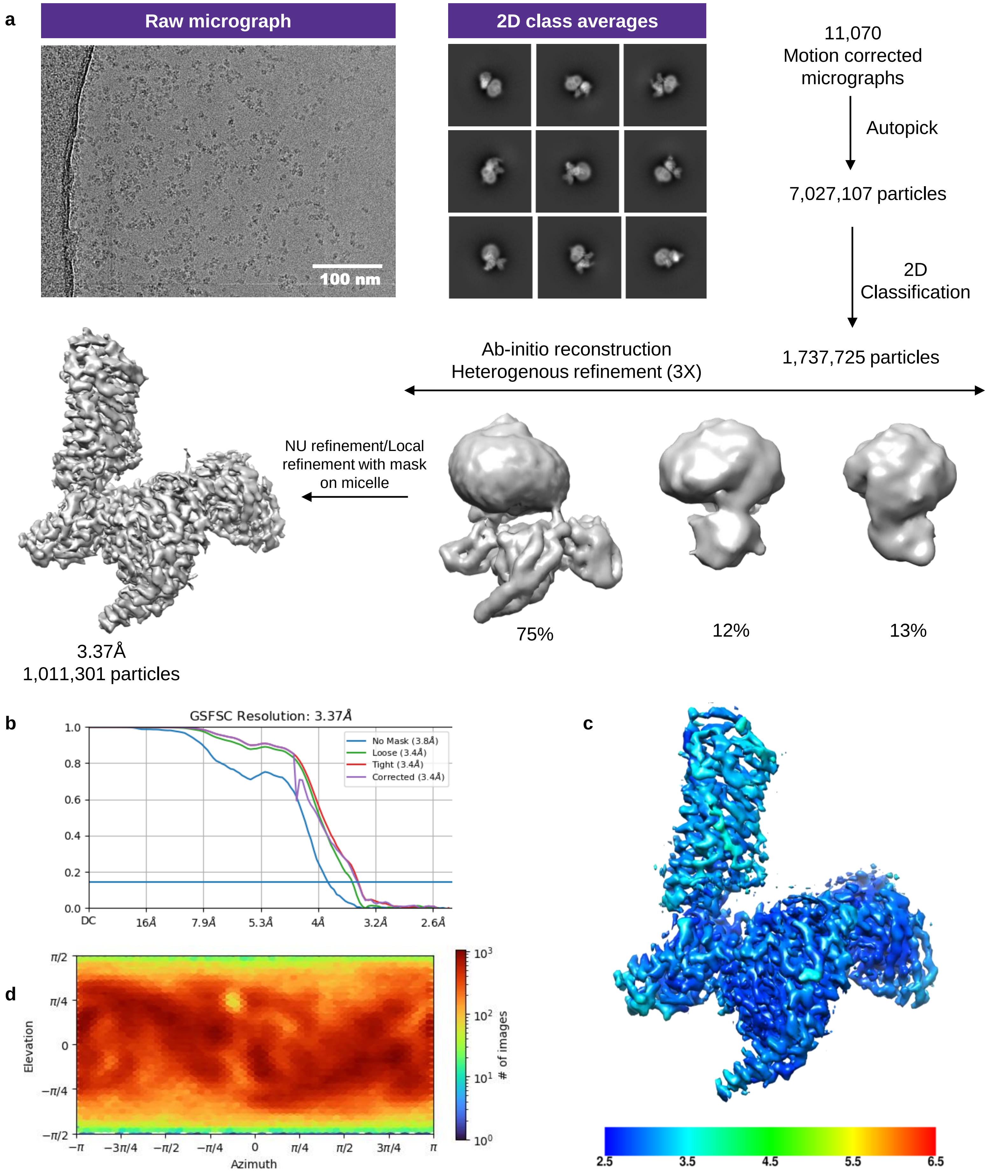


**Supplementary Figure 4: Cryo-EM data processing workflow of niacin-GPR109A-Go complex.**

**a,** Schematic representation of the cryo-EM data processing workflow. **b,** Gold standard fourier shell correlation curve (GFSC) at 0.143 threshold. **c,** Local resolution map of the 3D reconstruction (front view). **d,** Angular distribution of the particles used for final reconstruction.

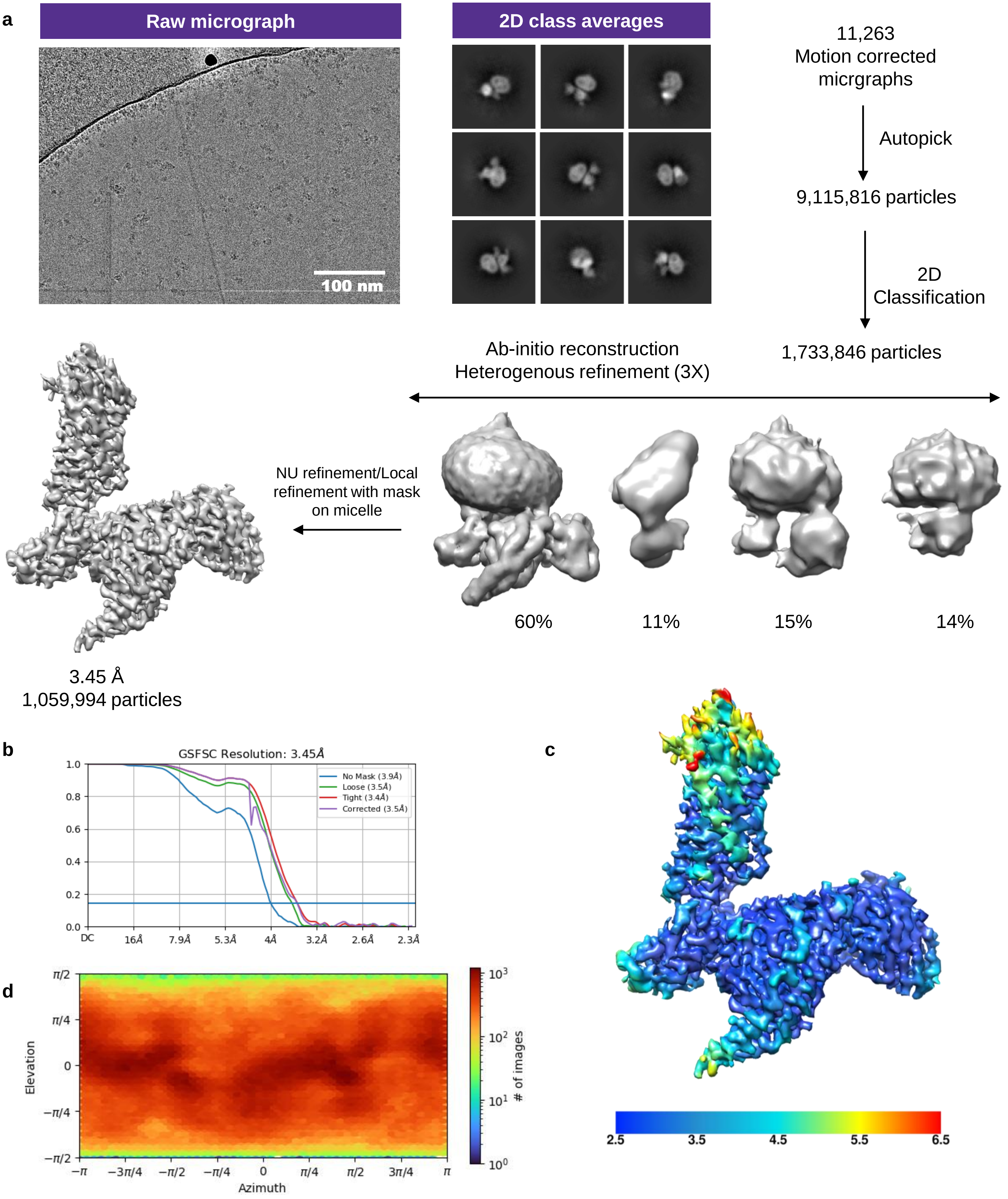


**Supplementary Figure 5: Data processing workflow of acipimox-GPR109A-Go complex.**

**a,** Flowchart of the cryo-EM data processing pipeline. **b,** Gold standard fourier shell correlation curve at 0.143 threshold. **c,** Local resolution map for the final 3D reconstruction (front view). **d,** Angular distribution of the particle set used for the final reconstruction.

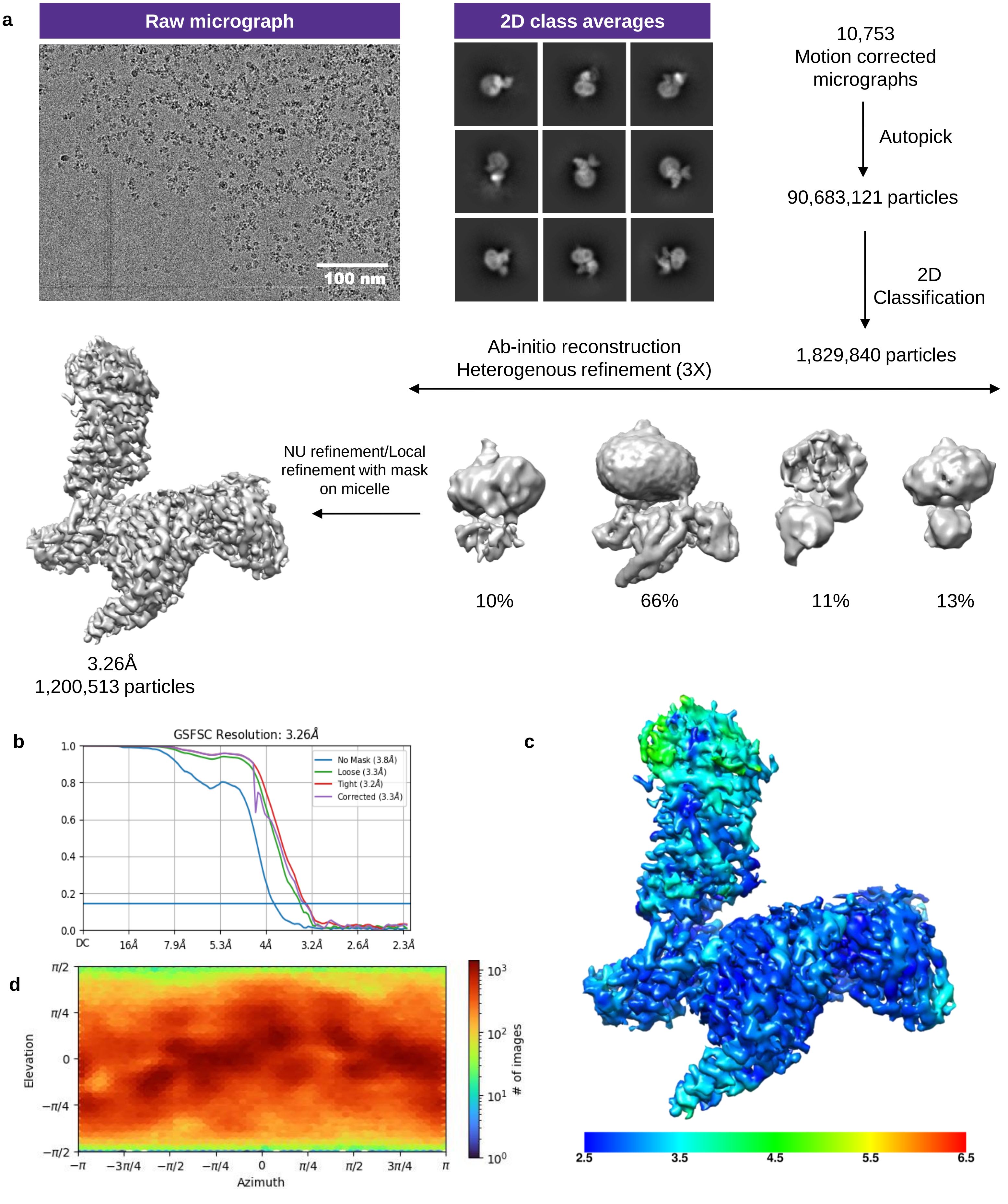


**Supplementary Figure 6: Cryo-EM reconstruction of MK6892-GPR109A-Go complex.**

**a,** Data processing workflow for reconstruction of MK6892-GPR109A-Go complex. **b,** Gold standard fourier shell correlation curve (GFSC) at a threshold of 0.143 indicates an overall resolution of 3.26Å. **c,** Local resolution map of the 3D reconstruction. **d,** Angular plot of the particles used for final 3D reconstruction of the complex.

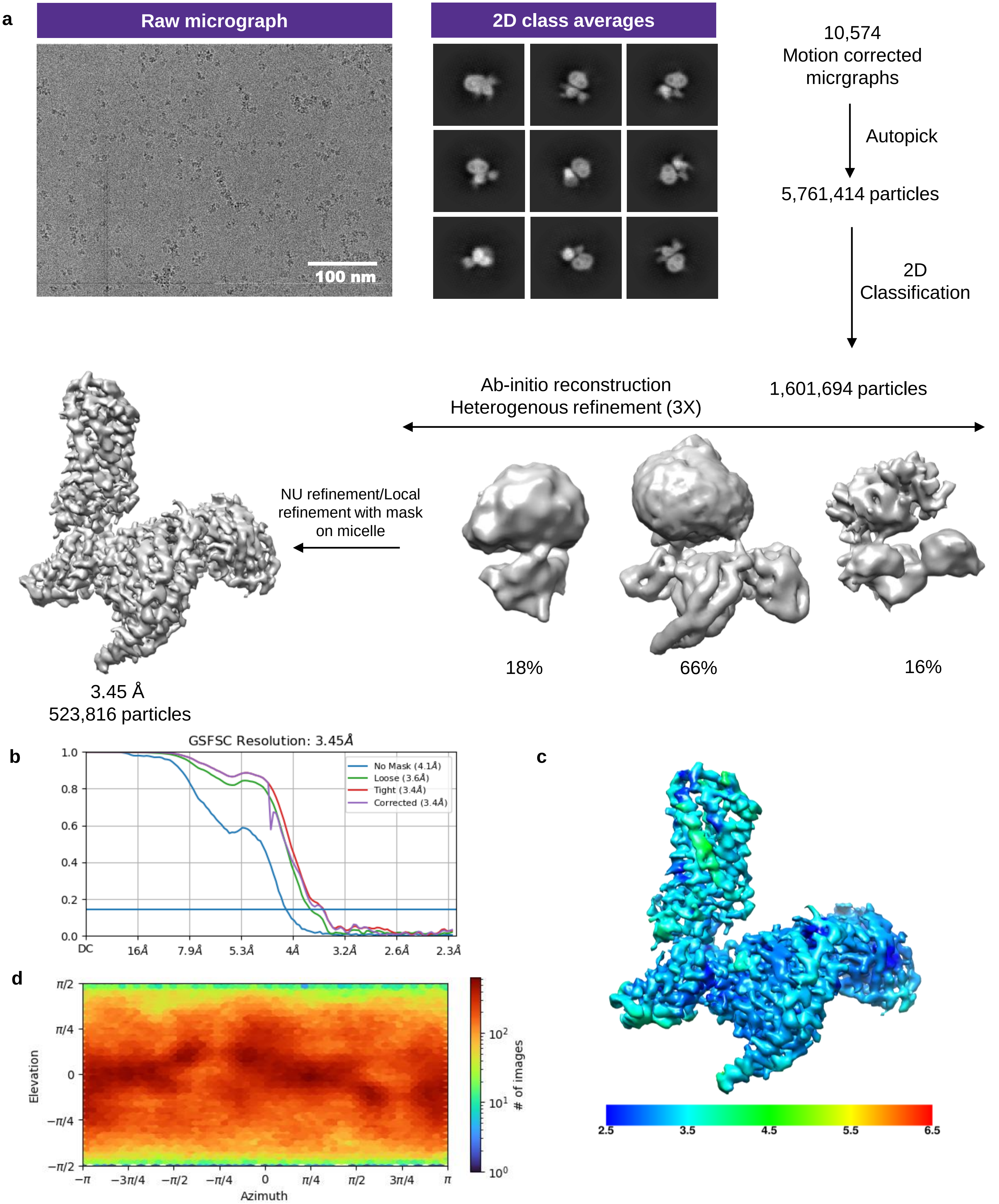


**Supplementary Figure 7: Data processing workflow of GSK256073-GPR109A-Go complex.**

**a,** Flowchart of the cryo-EM data processing pipeline. **b,** Gold standard fourier shell correlation curve at 0.143 threshold. **c,** Local resolution map for the final 3D reconstruction (front view). **d,** Angular distribution of the particle set used for the final reconstruction.

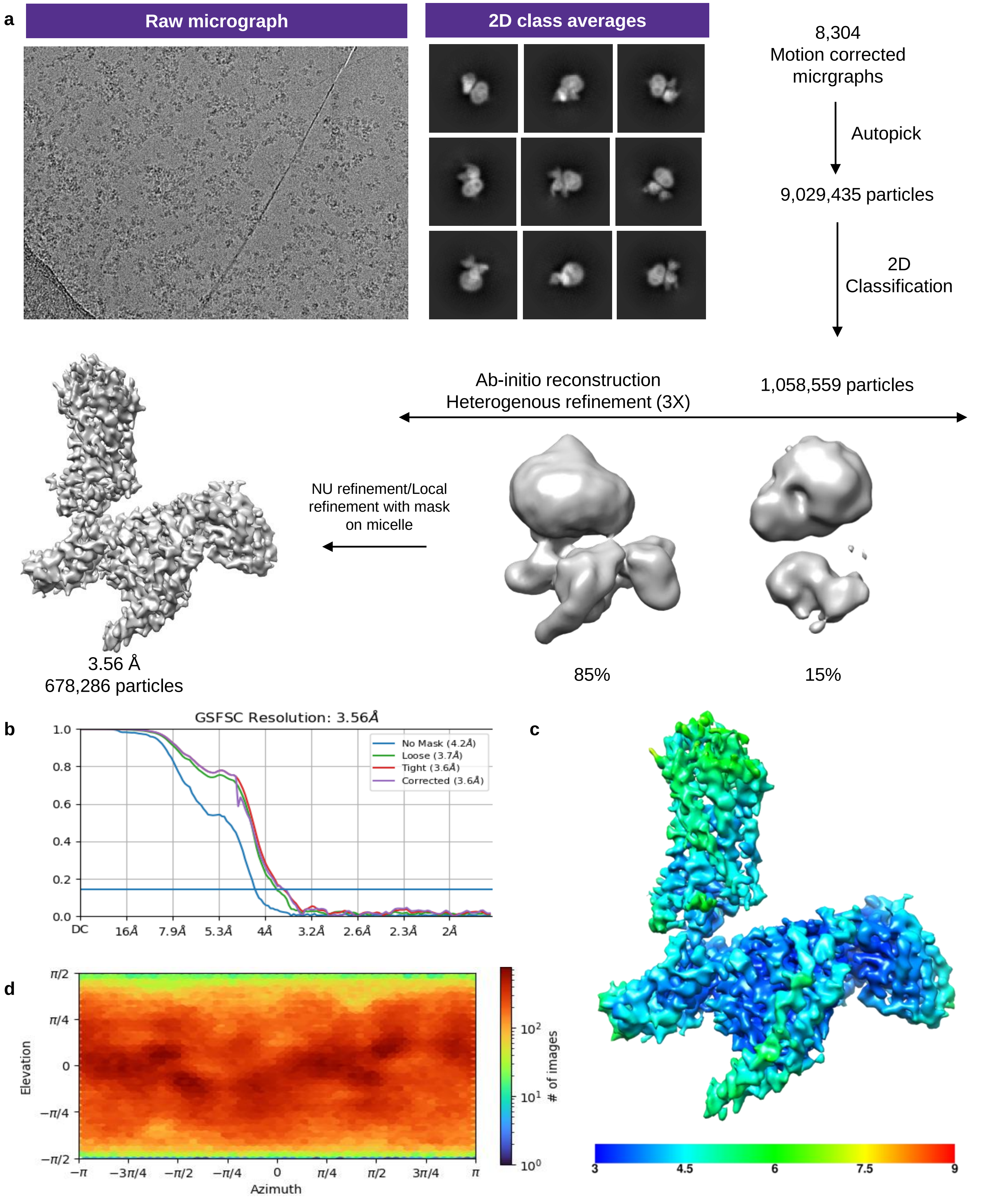


**Supplementary Figure 8: Data processing workflow of MMF-GPR109A-Go complex.**

**a,** Flowchart of the cryo-EM data processing pipeline. **b,** Gold standard fourier shell correlation curve at 0.143 threshold. **c,** Local resolution map for the final 3D reconstruction (front view). **d,** Angular distribution of the particle set used for the final reconstruction.

| Data collection and processing | | | | | |
|---|---|---|---|---|---|
| | Niacin-GPR109A-Go PDB 8IY9, EMD-35817 | Acipimox-GPR109A-Go PDB 8JER, EMD-36193 | GSK256073-GPR109A-Go PDB 8IYW, EMD-35831 | MK6892-GPR109A-Go PDB 8IYH, EMD-35822 | MMF-GPR109A-Go PDB 8JHN, EMD-36280 |
| Microscope | TFS Glacios | TFS Glacios | TFS Glacios | TFS Glacios | TFS Glacios |
| Camera | Gatan K3 | Gatan K3 | Gatan K3 | Gatan K3 | Gatan K3 |
| Magnification | 46,000x | 46,000x | 46,000x | 46,000x | 46,000x |
| Voltage (kV) | 200 | 200 | 200 | 200 | 200 |
| Defocus range (µm) | 0.5-2.5 | 0.5-2.5 | 0.5-2.5 | 0.5-2.5 | 0.5-2.5 |
| Exposure time (s) | 4 | 4 | 4 | 4 | 4 |
| Total dose ($e^-/A^2$) | 55 | 52 | 55 | 55 | 55 |
| Number of frames | 40 | 40 | 40 | 40 | 40 |
| Pixel size (Å) | 0.878 | 0.878 | 0.878 | 0.878 | 0.878 |
| Micrographs (no.) | 11,070 | 11,263 | 10,574 | 10,753 | 8,304 |
| Initial particles (no.) | 7,027,107 | 9,115,816 | 5,761,414 | 90,683,101 | 9,029,435 |
| Symmetry imposed | C1 | C1 | C1 | C1 | C1 |
| Final particles (no.) | 1,011,301 | 1,059,994 | 523,816 | 1,200,513 | 9,029,435 |
| Map resolution (Å) | 3.37 | 3.45 | 3.45 | 3.26 | 3.56 |
| FSC threshold | 0.143 | 0.143 | 0.143 | 0.143 | 0.143 |
| **Refinement** | | | | | |
| Initial model (PDB Code) | AlphaFold AF-Q8TDS4-F1 | 8IY9 | 8IY9 | 8IY9 | 8IY9 |
| Model resolution (Å) | 3.5 | 3.4 | 3.7 | 3.4 | 3.5 |
| FSC threshold | 0.5 | 0.5 | 0.5 | 0.5 | 0.5 |
| **Model composition** | | | | | |
| Non-hydrogen atoms | 8,371 | 8,322 | 8,322 | 8,386 | 8,376 |
| Protein residues | 1,133 | 1,132 | 1,136 | 1,133 | 1,133 |
| Ligand atoms | NIO=1 | ACI=1 | LIG=1 | FI7=1 | MMF=1 |
| **R.M.S. deviations** | | | | | |
| Bond length (Å) | 0.005 | 0.004 | 0.004 | 0.004 | 0.004 |
| Bond angle (º) | 0.895 | 1.011 | 0.748 | 1.034 | 1.008 |
| **Validation** | | | | | |
| Favored (%) | 97.13 | 96.86 | 97.14 | 97.67 | 97.31 |
| Allowed (%) | 2.87 | 3.14 | 2.86 | 2.33 | 2.69 |
| Disallowed (%) | 0 | 0 | 0 | 0 | 0 |
| MolProbity score | 1.40 | 1.63 | 1.48 | 1.35 | 1.60 |
| Clash Score | 4.33 | 8.16 | 5.94 | 5.27 | 8.66 |

**Supplementary Figure 9: Data collection, processing and refinement statistics of niacin, acipimox, MK6892, GSK256073 and MMF bound GPR109A-Go complexes.**

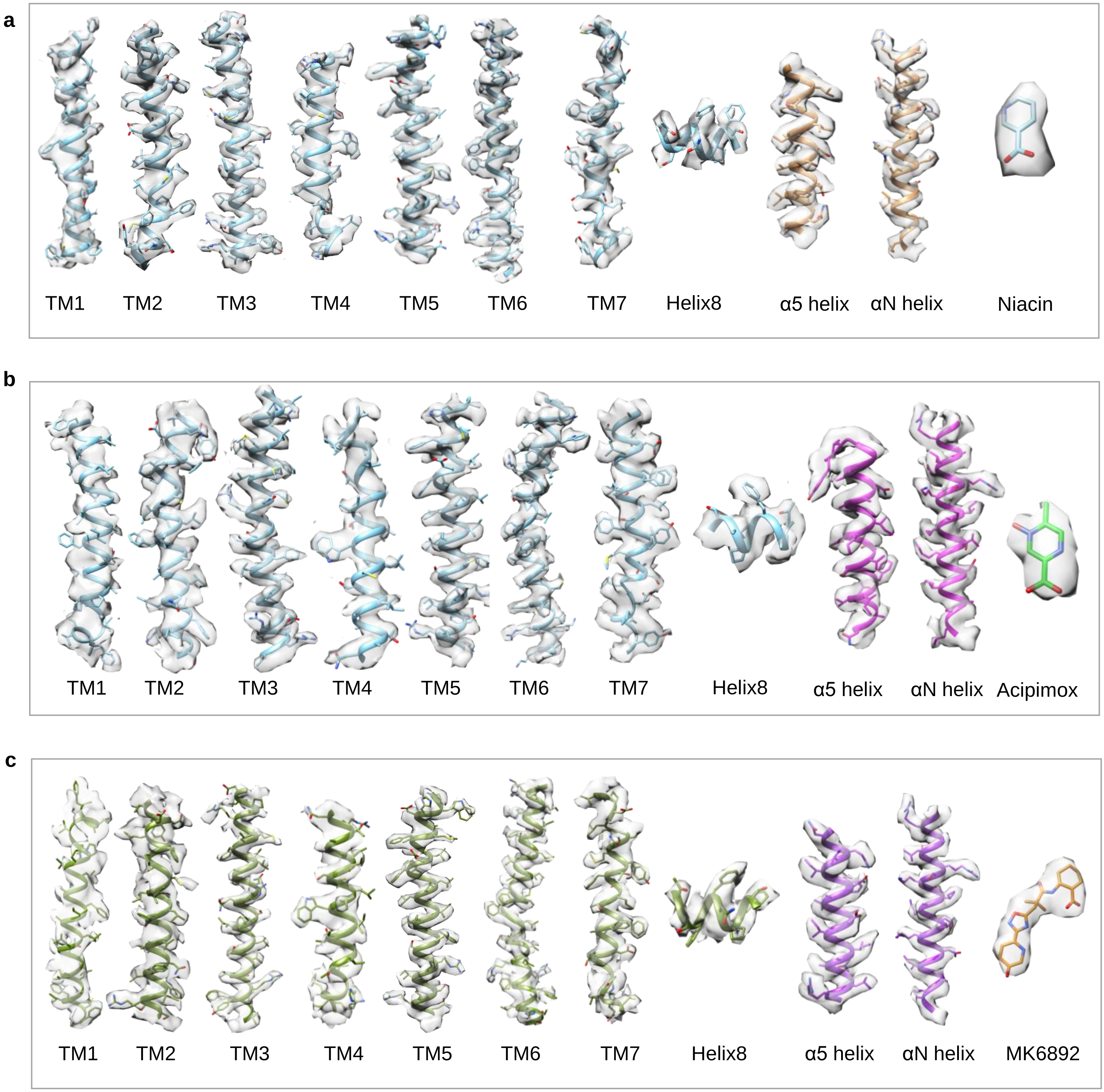
a
TM1
TM2
TM3
TM4
TM5
TM6
TM7
Helix8
α5 helix
αN helix
Niacin
b
TM1
TM2
TM3
TM4
TM5
TM6
TM7
Helix8
α5 helix
αN helix
Acipimox
c
TM1
TM2
TM3
TM4
TM5
TM6
TM7
Helix8
α5 helix
αN helix
MK6892

**d**

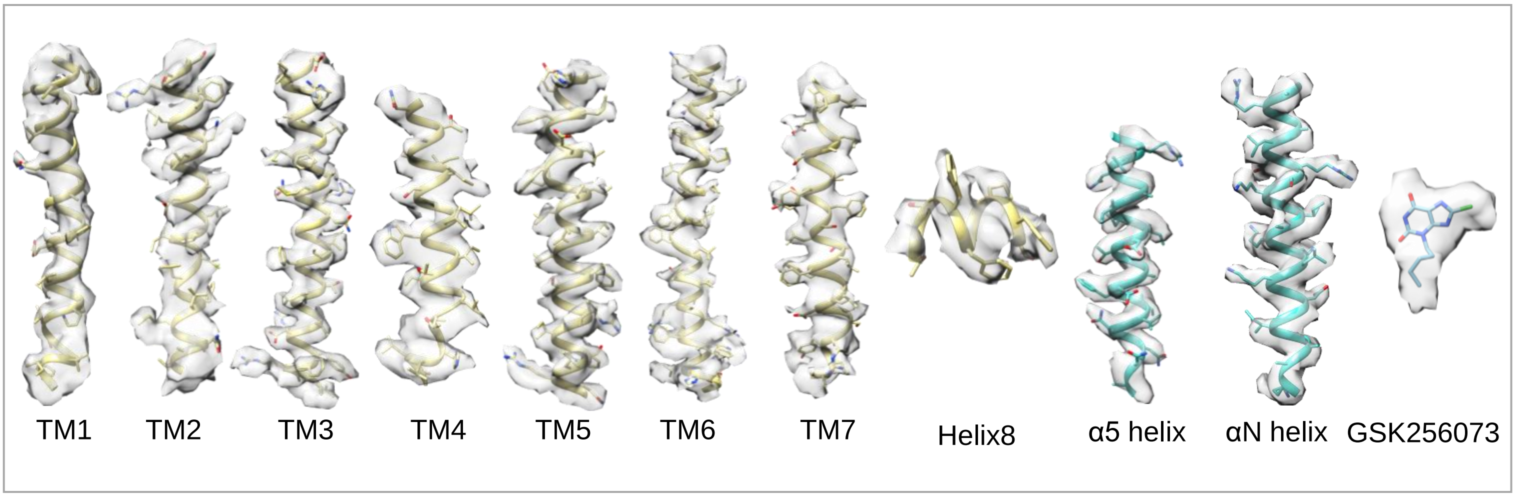


**e**

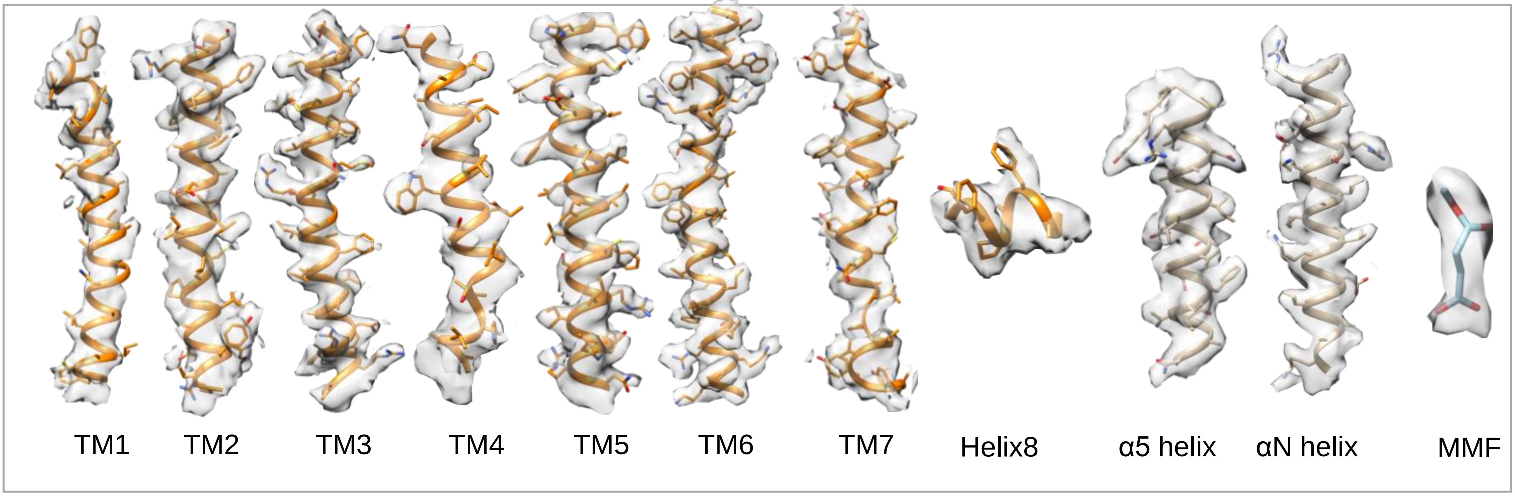


**Supplementary Figure 10: Representative electron density maps.**

**a,** EM densities for the TMs of niacin-GPR109A-Go structure (left to right): TM1 to TM7, helix 8, α5-helix, αN-helix, and niacin. **b,** EM densities for the TMs of Acipimox-GPR109A-Go structure (left to right): TM1 to TM7, α5-helix, αN-helix, helix 8 and acipimox **c,** EM densities for the TMs of MK6892-GPR109A-Go structure (left to right): TM1 to TM7, α5-helix, αN-helix, helix 8 and MK6892. **d,** EM densities for the TMs of GSK256073-GPR109A-Go structure (left to right): TM1 to TM7, α5-helix, αN-helix, helix 8 and GSK256073. **e,** EM densities for the TMs of MMF-GPR109A-Go structure (left to right): TM1 to TM7, α5-helix, αN-helix, helix 8 and MMF.

| Component | Total residues | Resolved residues | Total residues | Resolved residues |
|---|---|---|---|---|
| | Niacin-GPR109A-Go | | Acipimox-GPR109A-Go | |
| **Ligand/Drug** | 1 | 1 | 1 | 1 |
| **GPR109A** | M1-P363 | D8-T307 | M1-P363 | D8-T307 |
| **miniGαo** | M1-H57<br>T172-Y366 | S6-I55<br>T183-D230<br>H245-Y354 | M1-H57<br>T172-Y366 | S6-I55<br>T183-D230<br>H245-Y354 |
| **Gβ** | M1-N340 | E3-N340 | M1-N340 | E3-N340 |
| **Gγ** | M1-L71 | A7-R62 | M1-L71 | S8-R62 |
| **ScFv16** | D1-K248 | D1-R72<br>K76-S120<br>G135-K248 | D1-K248 | D1-R72<br>K76-S120<br>G135-K248 |

| Component | Total residues | Resolved residues | Total residues | Resolved residues |
|---|---|---|---|---|
| | MK6892-GPR109A-Go | | GSK256073-GPR109A-Go | |
| **Ligand/Drug** | 1 | 1 | 1 | 1 |
| **GPR109A** | M1-P363 | D8-T307 | M1-P363 | D8-T307 |
| **miniGαo** | M1-H57<br>T172-Y366 | S6-I55<br>T183-D230<br>H245-Y354 | M1-H57<br>T172-Y366 | S6-I155<br>T183-D230<br>H245-Y354 |
| **Gβ** | M1-N340 | E3-N340 | M1-N340 | E3-N340 |
| **Gγ** | M1-L71 | A7-R62 | M1-L71 | A7-R62 |
| **ScFv16** | D1-K248 | D1-R72<br>K76-S120<br>G135-K248 | D1-K248 | D1-S120<br>G135-K248 |

| Component | Total residues | Resolved residues |
|---|---|---|
| | MMF-GPR109A-Go | |
| **Ligand/Drug** | 1 | 1 |
| **GPR109A** | M1-P363 | D8-T307 |
| **miniGαo** | M1-H57<br>T172-Y366 | S6-I55<br>T183-D230<br>H245-Y354 |
| **Gβ** | M1-N340 | E3-N340 |
| **Gγ** | M1-L71 | A7-R62 |
| **ScFv16** | D1-K248 | D1-R72<br>K76-S120<br>G135-K248 |

**Supplementary Figure 11: Residues resolved in the structures of niacin, acipimox, MK6892, GSK256073 and MMF bound GPR109A-Go complexes.**

a

| Chain R (GPR109A) | Distance (Å) | Chain N (Niacin) |
|---|---|---|
| Leu83 (TM2) | 3.83 | Nio |
| Tyr87 (TM2) | 3.58 | Nio |
| Leu104 (TM3) | 3.65 | Nio |
| Leu107 (TM3) | 3.09 | Nio |
| Arg111 (TM3) | 2.66 | Nio |
| Ser178 (ECL2) | 3.69 | Nio |
| Ser179 (ECL2) | 3.38 | Nio |
| Phe180 (ECL2) | 3.64 | Nio |
| Phe277 (TM7) | 3.89 | Nio |
| Leu280 (TM7) | 3.64 | Nio |
| Tyr284 (TM7) | 2.56 | Nio |

b

| Chain R (GPR109A) | Distance (Å) | Chain Z (Acipimox) |
|---|---|---|
| Leu83 (TM2) | 3.84 | Aci401 |
| Tyr87 (TM2) | 3.05 | Aci401 |
| Leu104 (TM3) | 3.68 | Aci401 |
| Leu107 (TM3) | 3.25 | Aci401 |
| Arg111(TM3) | 2.34 | Aci401 |
| Cys177 (ECL2) | 3.72 | Aci401 |
| Ser178 (ECL2) | 3.18 | Aci401 |
| Ser179 (ECL2) | 2.54 | Aci401 |
| Phe180 (ECL2) | 3.31 | Aci401 |
| Phe277 (TM7) | 3.56 | Aci401 |
| Leu280 (TM7) | 3.51 | Aci401 |
| Tyr284 (TM7) | 2.72 | Aci401 |

**c**

| Chain D (GPR109A) | Distance (Å) | Chain R (MK6892) |
|---|---|---|
| Leu83 (TM2) | 3.68 | MK6892 |
| Tyr87 (TM2) | 3.04 | MK6892 |
| Leu104 (TM3) | 3.37 | MK6892 |
| Leu107 (TM3) | 2.88 | MK6892 |
| Ala108 (TM3) | 3.27 | MK6892 |
| Arg111 (TM3) | 3.38 | MK6892 |
| Gln112 (TM3) | 3.13 | MK6892 |
| Leu158 (TM4) | 2.79 | MK6892 |
| Thr159 (TM4) | 2.60 | MK6892 |
| Leu162 (TM4) | 3.26 | MK6892 |
| Ser179 (ECL2) | 2.40 | MK6892 |
| Phe180 (ECL2) | 3.19 | MK6892 |
| His189 (TM5) | 3.04 | MK6892 |
| Met192 (TM5) | 3.22 | MK6892 |
| Tyr284 (TM7) | 2.52 | MK6892 |

**d**

| Chain R (GPR109A) | Distance (Å) | Chain A (GSK256073) |
|---|---|---|
| Leu30 (TM1) | 3.85 | GSK256073 |
| Leu83 (TM2) | 3.03 | GSK256073 |
| Asn86 (TM2) | 3.13 | GSK256073 |
| Trp91 (ECL1) | 3.42 | GSK256073 |
| Leu107 (TM3) | 3.65 | GSK256073 |
| Arg111 (TM3) | 3.09 | GSK256073 |
| Ser178 (ECL2) | 3.49 | GSK256073 |
| Ser179 (ECL2) | 2.32 | GSK256073 |
| Phe180 (ECL2) | 2.60 | GSK256073 |
| Tyr284 (TM7) | 2.54 | GSK256073 |

**e**

| Chain R (GPR109A) | Distance (Å) | Chain Z (MMF) |
|---|---|---|
| Tyr87 (TM2) | 3.87 | MMF401 |
| Leu104 (TM3) | 3.60 | MMF401 |
| Leu107 (TM3) | 3.25 | MMF401 |
| Arg111 (TM3) | 2.71 | MMF401 |
| Ser178 (ECL2) | 3.20 | MMF401 |
| Ser179 (ECL2) | 2.67 | MMF401 |
| Phe180 (ECL2) | 3.48 | MMF401 |
| Phe277 (TM7) | 3.48 | MMF401 |
| Leu280 (TM7) | 3.42 | MMF401 |
| Tyr284 (TM7) | 2.47 | MMF401 |

**Supplementary Figure 12: (a, b, c, d, e) List of interaction between niacin, acipimox, MK6892, GSK256073 and MMF with GPR109A.**

**a**

| Chain R (GPR109A) | Distance (Å) | Chain A (Gαo) |
|---|---|---|
| Ser62 (TM2) | Gly350 (3.78), Cys351 (3.32) | Gly350, Cys351 |
| Arg63 (TM2) | Gly350 (3.81) | Gly350 |
| Arg125 (TM3) | Cys351 (3.28), Leu353 (3.52) | Cys351, Leu353 |
| Arg128 (TM3) | Asn347 (3.32) | Asn347 |
| Val129 (TM3) | Leu348 (3.53) | Leu348 |
| Pro132 (ICL2) | Ile343 (3.73), Ile344 (3.52) | Ile343, Ile344 |
| His133 (ICL2) | Leu195 (3.54), Thr340 (3.29) | Leu195, Thr340 |
| Arg218 (ICL3) | Thr340 (3.32), Asp341 (3.16), Ile344 (3.68) | Thr340, Asp341, Ile344 |
| Met220 (ICL3) | Ile344 (3.44) | Ile344 |
| His223 (TM6) | Glu318 (2.70) | Glu318 |
| Lys225 (TM6) | Tyr354 (2.97) | Tyr354 |
| Ile226 (TM6) | Tyr354 (3.58) | Tyr354 |
| Ala229 (TM6) | Leu353 (3.18) | Leu353 |
| Ile233 (TM6) | Leu353 (3.30) | Leu353 |
| Ser297 (TM7) | Gly352 (3.23) | Gly352 |
| Ser298 (Helix 8) | Gly352 (3.87) | Gly352 |
| Pro299 (Helix 8) | Tyr354 (3.86) | Tyr354 |

**b**

| Chain D (GPR109A) | Distance (Å) | Chain A (Gαo) |
|---|---|---|
| Lys60 (TM2) | Gly350 (3.28) | Gly350 |
| Ser62 (TM2) | Gly350 (2.88), Cys351 (3.39) | Gly350, Cys351 |
| Arg125 (TM3) | Leu353 (3.41) | Leu353 |
| Arg128 (TM3) | Asn347 (3.36), Cys351 (3.67) | Asn347, Cys351 |
| Val129 (TM3) | Leu348 (3.63) | Leu348 |
| Pro132 (ICL2) | Ile343 (3.88), Ile344 (3.49) | Ile344, Ile343 |
| His133 (ICL2) | Thr340 (2.75), Ile343(3.54) | Thr340, Ile343 |
| Arg218 (ICL3) | Asp337 (3.88), Thr340 (3.38), Asp341 (2.92) | Asp337, Thr340, Asp341 |
| Met220 (ICL3) | Ile344 (3.42) | Ile344 |
| His223 (TM6) | Glu318 (2.82) | Glu318 |
| Lys225 (TM6) | Tyr354 (3.48) | Tyr354 |
| Ile226 (TM6) | Tyr354 (3.54) | Tyr354 |
| Ala229 (TM6) | Leu353 (3.47) | Leu353 |
| Ile233 (TM6) | Leu353 (3.79) | Leu353 |
| Ser297 (TM7) | Leu353 (3.01), Tyr354 (3.65) | Leu353, Tyr354 |
| Ser298 (Helix 8) | Gly352 (3.37) | Gly352 |
| Pro299 (Helix 8) | Tyr354 (3.43) | Tyr354 |

c

| Chain R (GPR109A) | Distance (Å) | Chain A (GαO) |
|---|---|---|
| Lys60 (TM2) | Gly350 (3.53), Cys351 (3.88) | Gly350, Cys351 |
| Arg125 (TM3) | Cys351 (3.70), Leu353 (3.42) | Cys351, Leu353 |
| Arg128 (TM3) | Asn347 (3.48), Cys351 (3.21) | Asn347, Cys351 |
| Val129 (TM3) | Ile344 (3.73), Leu348 (3.77) | Ile344, Leu348 |
| Pro132 (ICL2) | Ile343 (3.65), Ile344 (3.62) | Ile343, Ile344 |
| His133 (ICL2) | Thr340 (3.28), Ile343 (3.64) | Thr340, Ile343 |
| Leu215 (TM5) | Ile344 (3.84) | Ile344 |
| Arg218 (ICL3) | Thr340 (3.31), Asp341 (3.23), Ile344 (3.77) | Thr340, Asp341, Ile344 |
| Met220 (ICL3) | Ile344 (3.58) | Ile344 |
| His223 (TM6) | Glu318 (3.03) | Glu318 |
| Lys225 (TM6) | Tyr354 (3.39) | Tyr354 |
| Ile226 (TM6) | Tyr354 (3.57) | Tyr354 |
| Arg228 (TM6) | | |
| Ala229 (TM6) | Leu353 (3.50) | Leu353 |
| Ser297 (TM7) | Gly352 (3.12) | Gly352 |
| Ser298 (H8) | Gly352 (2.91), Leu353 (3.09), Tyr354 (3.49) | Gly352, Leu353, Tyr354 |

d

| Chain R (GPR109A) | Distance (Å) | Chain B (Gαo) |
|---|---|---|
| Arg125 (TM3) | Leu353 (3.40) | Leu353 |
| Arg128 (TM3) | Asn347 (3.01), Gly350 (3.65), Cys351 (3.47) | Asn347, Gly350, Cys351 |
| Val129 (TM3) | Ile344 (3.84), Leu348 (3.57) | Ile344, Leu348 |
| Pro132 (ICL2) | Ile344 (3.64), Asn347 (3.39) | Ile344, Asn347 |
| His133 (ICL2) | Leu195 (3.86), Thr340 (3.39) | Leu195, Thr340 |
| Lys138 (ICL2) | Ala31 (3.80) | Ala31 |
| Arg218 (ICL3) | Thr340 (3.31), Asp341 (3.10), Ile344 (3.71) | Thr340, Asp341, Ile344 |
| Met220 (ICL3) | Ile344 (3.52) | Ile344 |
| His223 (TM6) | Glu318 (2.55) | Glu318 |
| Lys225 (TM6) | Tyr354 (3.49), | Tyr354 |
| Ile226 (TM6) | Leu348 (3.80), Tyr354 (3.42) | Leu348, Tyr354 |
| Ala229 (TM6) | Leu353 (3.39) | Leu353 |
| Ile233 (TM6) | Leu353 (3.72) | Leu353 |
| Ser298 (Helix 8) | Leu353 (3.39) | Leu353 |

**e**

| Chain R (GPR109A) | Distance (Å) | Chain A (Gαo) |
|---|---|---|
| Lys60 (TM2) | Gly350 (3.24) | Gly350 |
| Ser62 (TM2) | Gly350 (3.51), Cys351 (3.75) | Gly350, Cys351 |
| Asp124 (TM3) | Cys351 (3.56) | Cys351 |
| Arg125 (TM3) | Leu353 (3.40) | Leu353 |
| Arg128 (TM3) | Asn347 (3.34), Cys351 (2.78) | Asn347, Cys351 |
| Val129 (TM3) | Leu348 (3.55) | Leu348 |
| Pro132 (ICL2) | Ile344 (3.60) | Ile344 |
| His133 (ICL2) | Leu195 (3.51), Thr340 (3.26), Ile343 (3.58) | Leu195, Thr340, Ile343 |
| Leu215 (TM5) | Ile344 (3.78) | Ile344 |
| Arg218 (ICL3) | Thr340 (3.36), Asp341 (3.22), Ile344 (3.85) | Thr340, Asp341, Ile344 |
| Met220 (ICL3) | Ile344 (3.47) | Ile344 |
| His223 (TM6) | Glu318 (2.76) | Glu318 |
| Lys225 (TM6) | Tyr354 (3.22) | Tyr354 |
| Ile226 (TM6) | Tyr354 (3.52) | Tyr354 |
| Arg228 (TM6) | Tyr354 (3.81) | Tyr354 |
| Ala229 (TM6) | Leu353 (3.48) | Leu353 |

**Supplementary Figure 13: (a, b, c, d, e) List of GPR109A-Go interactions in the structures of niacin, acipimox, MK6892, GSK256073 and MMF-GPR109A-Go.**

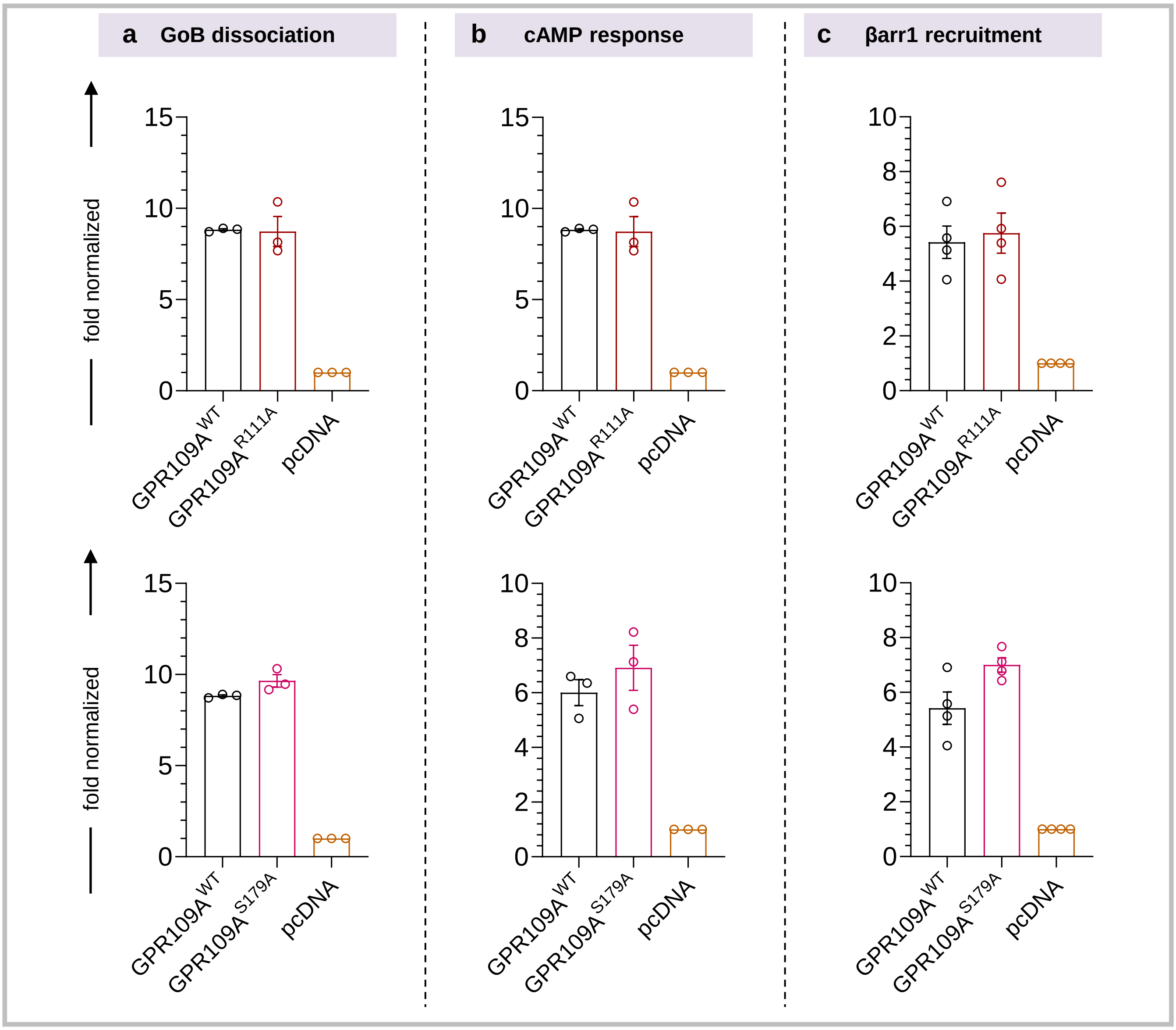


**Supplementary Figure 14 : Surface expression of GPR109A$^{WT}$ and mutants in various assays.**

**a,** Surface expression of GPR109A$^{WT}$, GPR109A$^{R111A}$ , and GPR109A$^{S179A}$ in GoB dissociation assay was measured using whole cell based surface ELISA (mean±SEM; n=3; normalized as fold over pcDNA) **b,** GPR109A$^{WT}$ and mutants surface expression in GloSensor assay (mean±SEM; n=4; normalized as fold over pcDNA) **c,** Surface expression of GPR109A$^{WT}$ and mutants in β-arrestin recruitment assay (mean±SEM; n=4; normalized as fold over pcDNA).
